# Localized co-inoculation of *Bacillus subtilis* and *Trichoderma afroharzianum* acts synergistically to reshape the root microbiome and improve plant performance in sorghum

**DOI:** 10.64898/2026.08.11.744214

**Authors:** Bishrant Pant, Maruf Khan, Ahmad H. Kabir

## Abstract

Despite their agricultural potential, how bacterial–fungal consortia reshape root microbiomes and improve crop performance in sorghum remains poorly understood. Here, we investigated how individual and combined inoculation with *Bacillus subtilis* and *Trichoderma afroharzianum* influenced sorghum performance and root microbiome assembly. The in vitro co-culture assay demonstrated the compatibility of *B. subtilis* and *T. afroharzianum* as a microbial consortium. The *B. subtilis*–*T. afroharzianum* consortium demonstrated the highest CPPI (composite plant performance index) and shoot fresh weight in sorghum, while all inoculation treatments improved multiple growth and physiological traits. Split-root analysis demonstrated that bilateral root co-inoculation was necessary to maximize whole-plant growth benefits. Also, *B. subtilis*–*T. afroharzianum* co-inoculation increased carbon levels in both roots and leaves, accompanied by enhanced rhizosphere siderophore production consistent with improved nutrient status. In microbial community analysis, neither bacterial nor fungal alpha or beta diversity differed significantly among treatments; instead, inoculation selectively restructured root microbial communities. The *B. subtilis–T. afroharzianum* consortium selectively enriched plant growth-promoting *Actinoplanes*, siderophore-producing *Enterobacter*, and the plant-beneficial fungal genus *Podospora*. Co-occurrence network analysis identified *Rhodoplanes, Serendipita,* and *Zopfiella* among hub taxa associated with *B. subtilis–T. afroharzianum* co-inoculation, suggesting potential roles in microbial community connectivity and organization. Furthermore, the persistence of *Streptomyces* and *Serendipita*, particularly the latter, suggests the presence of a beneficial microbial core that may contribute to sustained rhizosphere functioning. In addition, *Bacillus* and *Serendipita* were among the indicator taxa associated with inoculated treatment combinations, suggesting that the inoculants selectively assembled a distinct plant-beneficial microbiome. *Devosia* was associated with chlorophyll content, siderophore production, and shoot height, whereas *Serendipita* correlated with shoot biomass under the *B. subtilis*–*T. afroharzianum* co-inoculation. Taken together, *B. subtilis–T. afroharzianum* consortium promotes sorghum growth by selectively reshaping the root microbiome, highlighting its potential as a next-generation microbial biofertilizer.

## 1. Introduction

Sorghum (*Sorghum bicolor* L. Moench) is the fifth most important cereal crop worldwide and is cultivated for grain, forage, and bioenergy because of its high biomass productivity, efficient water use, and adaptability to diverse environmental conditions (Rooney et al. 2007). Improving sorghum productivity using sustainable biological inputs has become increasingly important as agriculture seeks alternatives to heavy reliance on synthetic fertilizers and chemical amendments. Plant-associated microorganisms, especially rhizosphere bacteria and fungi, can improve crop performance by improving nutrient acquisition, phytohormone regulation, root development, photosynthetic activity, and stress resilience (Compant et al., 2019; de Vries et al., 2020). In sorghum, microbial inoculation has been shown to influence root architecture, biomass accumulation, and rhizosphere microbial assembly, highlighting the importance of beneficial plant–microbe interactions in this crop (Sahib et al., 2020).

Among beneficial microbial inoculants, *Bacillus subtilis* is one of the most widely studied plant-growth-promoting rhizobacteria because of its strong rhizosphere competence, spore-forming ability, and multiple growth-promoting mechanisms. These include nutrient solubilization, production of phytohormones and volatile compounds, pathogen suppression, biofilm formation, and modulation of plant stress responses (Hashem et al., 2019; Blake et al., 2021; Mahapatra et al., 2022). Similarly, *Trichoderma* species, including *Trichoderma afroharzianum*, are recognized as beneficial fungi capable of promoting plant growth through root colonization, secretion of bioactive metabolites, improvement of nutrient availability, and activation of plant defense and stress-responsive pathways (Contreras-Cornejo et al., 2024). Furthermore, beneficial root-associated microorganisms can influence plant performance through both local and systemic signaling mechanisms. Local perception of microbial signals at colonized root regions can modify root architecture and symbiotic signaling, while microbe-induced signals can also be transmitted to distal, non-colonized tissues, resulting in systemic physiological responses (Kabir et al., 2024; Saiz-Fernández et al., 2021). Recent work further indicates that *T. afroharzianum* T22 can alter endophytic microbial communities associated with growth promotion in sorghum (Reid et al., 2024).

Although individual microbial inoculants such as *B. subtilis* and *T. afroharzianum* have consistently promoted plant growth, increasing evidence suggests that microbial consortia often outperform single inoculants because different microorganisms can provide complementary functions within the rhizosphere. Bacterial and fungal partners may occupy distinct ecological niches and interact synergistically to enhance nutrient mobilization, phytohormone production, pathogen suppression, and rhizosphere colonization, thereby improving plant growth more consistently across diverse environmental conditions (Berg et al., 2020; Gastélum et al., 2025). Several studies have reported that combined inoculation of plant growth-promoting bacteria and fungi results in greater biomass accumulation, nutrient uptake, and stress tolerance than either microorganism alone, although the magnitude of these benefits depends on the compatibility of the microbial partners and their interactions with the native microbiome (Timmusk et al., 2017; Jacoby et al., 2017).

Despite the importance of microbial inoculants, little is known about how combined inoculation with *B. subtilis* and *T. afroharzianum* influences bacterial and fungal community assembly, microbial interaction networks, and plant–microbiome relationships in sorghum. Addressing these knowledge gaps will advance our understanding of microbiome-assisted crop production and facilitate the rational development of effective microbial consortia for sustainable agriculture. Therefore, we combined plant phenotyping with amplicon sequencing, microbial network analysis, core microbiome analysis, indicator species analysis, and plant–microbe correlation analysis to investigate how individual and combined inoculation with *B. subtilis* and *T. afroharzianum* influences sorghum performance and root microbiome assembly. We also integrated a Composite Plant Performance Index (CPPI) with treatment-specific plant–microbiome correlation analyses to link microbial community shifts with overall plant performance.

## 2. Materials and methods

### 2.1. Dual-culture compatibility assay

The compatibility between *T. afroharzianum* and *B. subtilis* was evaluated using a dual-culture assay on half-strength potato dextrose agar (½ PDA; Difco Laboratories, Detroit, MI, USA). A 5-mm-diameter agar plug was excised from the actively growing margin of a 5-day-old *T. afroharzianum* culture and placed approximately 1 cm from one edge of a 90-mm Petri dish. *B. subtilis* was adjusted to approximately 1 × 10^8 CFU mL ¹ (OD600 ≈ 0.1) and inoculated as a single linear streak on the opposite side of the plate, approximately 5 cm from the fungal plug. Control plates containing only *T. afroharzianum* or *B. subtilis* were prepared similarly. All plates were incubated at 28 ± 1 °C in the dark. Fungal colony diameter was measured along the axis toward the bacterial streak, whereas bacterial colony width was measured perpendicular to the streak at 48, 72, and 96 h after inoculation. Each treatment consisted of five independent biological replicates (*n* = 5). Colony morphology, sporulation, and the presence of inhibition zones were visually assessed throughout the incubation period.

### 2.2. Plant cultivation and growth conditions

Seeds of sorghum (*Sorghum bicolor* L.; USDA-GRIN accession PI 591038, sweet sorghum) were surface-sterilized with 2% sodium hypochlorite for 5 min, rinsed three times with sterile distilled water, and germinated in seedling trays at 25 °C for 48 h. Uniform seedlings were transplanted into plastic pots containing 500 g of a soil substrate consisting of agricultural field soil and commercial potting mix (1:2, v/v). Four treatments were established: (i) uninoculated control, (ii) *B. subtilis* (BS; 1 mL of 1 × 10 CFU mL ¹; ARS Culture Collection–USDA, NRRL B-14596), (iii) *T. afroharzianum* T22 (TA; 1 mL of 1 × 10 conidia mL ¹), and (iv) combined BS + TA inoculation. Each treatment consisted of three biological replicates, with one plant grown per pot. Microbial inoculants were applied as a soil drench at the time of seedling transplantation. Soil pH was maintained at approximately 6.5 throughout the experiment. Plants were grown for six weeks in a greenhouse under a completely randomized design with a 10-h light/14-h dark photoperiod, a photosynthetic photon flux density of approximately 250 μmol m ² s ¹, and a temperature of 25 ± 2 °C.

### 2.3. Split root experiments

A split-root experiment was conducted to determine whether the growth-promoting effects of *B. subtilis* (BS) and *T. afroharzianum* (TA) consortium depended on local root colonization or could be transmitted to the non-inoculated root compartment. Briefly, sorghum seedlings were initially grown in sterile soil for 2 weeks. Following transfer, plants were maintained in the split-root system for an additional four weeks under the greenhouse conditions described above. The root system of each seedling was then carefully divided into two approximately equal portions and transplanted into two adjacent soil compartments (Fig. 4A). Plants were assigned to three treatments: SR1, control/control, in which neither root compartment received microbial inoculum; SR2, control/BS+TA, in which only one root compartment was inoculated with the BS+TA consortium; and SR3, BS+TA/BS+TA, in which both root compartments were inoculated. Each treatment consisted of three independent biological replicates.

### 2.4. Molecular detection of inoculants in root samples

Microbial colonization of sorghum roots by TA and BS was verified by PCR. Root samples were gently rinsed twice with sterile phosphate-buffered saline (PBS), briefly vortexed to remove loosely attached soil particles, and washed twice with sterile distilled water. Genomic DNA was extracted from approximately 0.2 g of root tissue using the cetyltrimethylammonium bromide (CTAB) method (Clarke, 2009). DNA concentration and purity were determined using a NanoDrop ND-1000 spectrophotometer (Thermo Fisher Scientific, Wilmington, DE, USA), and DNA samples were normalized to equal concentrations prior to PCR. PCR amplification was performed using TA-specific primers (forward: 5′-GTCGGTAGCTGAAAGGGG-3′; reverse: 5′-ATTAGGCCGGAAACACC-3′) and BS-specific primers (forward: 5′-TCTGCTCGTGAACGGTGCT-3′; reverse: 5′-TTTCGCCTTATTTACTTGG-3′) (IARRPCAAS, 2013). Each PCR reaction (25 μL) contained 1× PCR buffer, 0.2 mM dNTPs, 0.4 μM of each primer, 1 U Taq DNA polymerase, and approximately 50 ng of genomic DNA. The amplification conditions consisted of an initial denaturation at 95 °C for 3 min, followed by 35 cycles of 95 °C for 30 s, primer-specific annealing at 56 °C for 30 s, and 72 °C for 30 s, with a final extension at 72 °C for 5 min. PCR products were separated by electrophoresis on a 1.5% agarose gel stained with GelRed and visualized using a GelDoc™ EZ Imager (Bio-Rad Laboratories, Hercules, CA, USA).

### 2.5. Growth and physiological measurements

Chlorophyll content was determined non-destructively using a SPAD chlorophyll meter (SPAD-502 Plus, Konica Minolta, Japan). Chlorophyll fluorescence kinetics (OJIP), including the maximum quantum efficiency of PSII (Fv/Fm) and the photosynthetic performance index (Pi_ABS), were measured on the uppermost fully expanded leaves at three different positions using a portable FluorPen FP 110 (Photon Systems Instruments, Czech Republic). Before OJIP measurements, leaves were dark-adapted for 1 h. Furthermore, shoot height was measured from the stem base to the tip of the longest leaf using a ruler, whereas root length was measured from the crown to the tip of the longest root after gently washing the root system. After morphological measurements, shoots and roots were separated and immediately weighed to determine shoot fresh weight and root fresh weight using an analytical balance.

### 2.6. Composite plant performance index

The Composite Plant Performance Index (CPPI) was calculated to integrate the measured growth and physiological traits into a single performance metric. For each trait, treatment means were standardized using Z-scores, calculated as Z=(x−μ)/σ, where x is the mean, μ is the overall mean across treatments, and σ is the corresponding standard deviation. The CPPI for each treatment was obtained by averaging the Z-scores of all included traits, with higher values indicating better overall plant performance.

### 2.7. Elemental analysis in sorghum root and leaf

Root samples were carefully harvested and thoroughly rinsed under running tap water to remove adhering soil particles, followed by immersion in 0.1 mM CaSO_4_ solution for 10 min to eliminate surface-bound ions. The roots were then rinsed with deionized water, whereas leaf samples were washed exclusively with deionized water. All samples were transferred to labelled paper envelopes and dried in a forced-air oven at 75 °C for 72 h until a constant weight was obtained. The dried root and leaf tissues were subsequently ground into a fine powder for elemental analysis. Iron (Fe) concentration was determined using inductively coupled plasma optical emission spectrometry (ICP-OES; Shimadzu ICPE-9820, Japan), whereas nitrogen (N) and carbon (C) contents were quantified using an elemental analyzer (Thermo Fisher FLASH SMART, USA). All analyses were conducted at the Center for Advances in Water and Air Quality, Lamar University.

### 2.8. Rhizosphere siderophore assay

Rhizosphere soil was collected by gently brushing soil adhering to the roots. Siderophore production was quantified using the Chrome Azurol S (CAS) assay (Alexander & Zuberer, 1991). Briefly, rhizosphere soil suspensions were prepared in sterile distilled water, and the supernatant was mixed with CAS reagent. After incubation at room temperature, absorbance was measured at 630 nm using a spectrophotometer. Siderophore production was expressed as percent siderophore using the formula: siderophore (%) = [(Ar − As)/Ar] × 100, where Ar is the absorbance of the CAS reagent (reference) and As is the absorbance of the sample.

### 2.9. Amplicon sequencing and data processing

Root microbial communities were characterized using Illumina amplicon sequencing by targeting the bacterial 16S rRNA gene and the fungal internal transcribed spacer (ITS) region. Amplicon libraries were prepared by amplifying the bacterial 16S rRNA (V3V4) gene using the primer pair 338F (5′- ACTCCTACGGGAGGCAGCAG-3′) and 806R (5′-GGACTACHVGGGTWTCTAAT-3′), and the fungal internal transcribed spacer (ITS) region using the primer pair ITS1FI2 (5′- GAACCWGCGGARGGATCA-3′) and ITS2 (5′-GCTGCGTTCTTCATCGATGC-3′). Paired-end sequencing (2 × 250 bp) was performed on the Illumina NovaSeq 6000 platform.

Raw sequence reads were quality-filtered and adapter-trimmed using Cutadapt (Martin, 2011), followed by denoising, chimera removal, and inference of amplicon sequence variants (ASVs) using the DADA2 pipeline (Callahan et al., 2016). Reads assigned to chloroplast and mitochondrial DNA were excluded from the bacterial dataset before downstream analyses. Taxonomic assignment of bacterial ASVs (amplicon sequence variants) was performed against the SILVA 138 reference database (Quast et al., 2013), whereas fungal ASVs were classified using the UNITE database (Nilsson et al., 2019). All downstream analyses were performed in R using the phyloseq (McMurdie & Holmes, 2013) and related packages. Alpha diversity was estimated using the observed ASVs and Shannon diversity indices after rarefaction to an even sequencing depth. Beta diversity was assessed using non-metric multidimensional scaling (NMDS) based on Bray–Curtis dissimilarity. Community differences between treatments were tested using permutational multivariate analysis of variance (PERMANOVA). The relative abundances of bacterial taxa were calculated after agglomerating ASVs at the phylum, class, order, family, and genus levels using the phyloseq package, and the top 15 most abundant taxa at each taxonomic rank were visualized as stacked bar plots based on relative abundance. Differences in the relative abundance of each taxon among treatments were assessed using the non-parametric Kruskal–Wallis test, with significance determined at *P* < 0.05. Further, differential abundance analysis was performed using DESeq2 after converting the phyloseq object to a DESeq2 dataset with treatment as the design factor. Pairwise comparisons were conducted among treatments. ASVs were considered significantly enriched or depleted at an adjusted P < 0.05 and |log fold change| > 1, and the results were visualized using volcano plots.

### 2.10. Global co-occurrence network analysis

Bacterial and fungal ASVs were separately agglomerated to the genus level using the phyloseq package, and genus abundances were transformed to relative abundance. Genera with a mean relative abundance <0.1% or zero variance across samples were excluded before analysis. A global microbial co-occurrence network was constructed using pairwise Spearman’s rank correlations among retained genera. P-values were adjusted for multiple testing using the Benjamini–Hochberg procedure, and correlations with |ρ| ≥ 0.60 and false-discovery-rate-adjusted P < 0.05 were retained to generate an undirected network in igraph, and isolated nodes were removed. Hub taxa were identified based on degree centrality, with additional centrality metrics (betweenness, closeness, and eigenvector centrality) calculated to characterize node importance. To associate hub taxa with experimental treatments, the mean relative abundance of each genus was calculated for each treatment, and each hub taxon was assigned to the treatment in which it exhibited the highest average abundance. The top hub taxa overall and the top five treatment-associated hub taxa were visualized according to degree centrality.

### 2.11. Core microbiome and indicator species analysis

Core microbiomes were identified separately for bacterial and fungal communities at the genus level. Genera present in at least 70% of samples with a mean relative abundance ≥0.1% were defined as core taxa. Relative abundances of the core genera were calculated, and the ten most abundant genera were compared among the treatment groups. Furthermore, indicator species analysis was performed at the genus level in R using the indicspecies package. Relative abundance data were used to identify microbial taxa significantly associated with each treatment or host plant. Indicator values (IndVal) were calculated based on the specificity and fidelity of each genus to a given group, and statistical significance was assessed using 999 permutation tests. The top indicator taxa with significant (*P* < 0.05) indicator values were visualized as horizontal bar plots using ggplot2.

### 2.12. Correlation analysis between microbial genera and plant traits

Correlation analysis was performed in R using Spearman’s rank correlation to evaluate associations between bacterial and fungal genera and plant growth and physiological traits. Genus-level relative-abundance data were used, and the 30 most abundant genera, based on mean relative abundance across all samples, were included in the analysis. Pairwise correlations were calculated between genus-level relative abundances and measured plant traits. Correlations with *P* < 0.05 were considered statistically significant. Correlation coefficients were visualized as clustered heatmaps using the pheatmap package in R, with hierarchical clustering applied to both genera and plant traits based on correlation similarity. Positive and negative correlations are represented by red and blue colors, respectively.

### 2.13. Statistical analysis

Growth and physiological experimental data were analyzed using R software (version 4.4.1). Differences among treatments were determined by one-way analysis of variance (ANOVA) with treatment as the fixed effect. When significant differences were detected (*P* < 0.05), treatment means were compared using Fisher’s least significant difference (LSD) test, and different lowercase letters indicate significant differences among treatments. Data are presented as mean ± standard deviation (SD) of three independent biological replicates (*n* = 3).

## 3. Results

### 3.1. In vitro compatibility between T. afroharzianum and B. subtilis

The compatibility between *T. afroharzianum* and *B. subtilis* was evaluated using a dual-culture assay on half-strength PDA medium. Throughout the 96-h incubation period, no visible inhibition zone developed between the fungal colony and the bacterial streak (Fig. 1A). Similarly, neither microorganism exhibited detectable changes in colony morphology, pigmentation, or sporulation in the presence of the other, indicating that the two microorganisms were mutually compatible under the tested conditions (Fig. 1A). Quantitative measurements further supported this observation (Fig. 1B). The growth of *B. subtilis* in co-culture was comparable to that in single culture at 48, 72, and 96 h, with no significant differences detected at any time point. Similarly, colony expansion of *T. afroharzianum* was unaffected by co-cultivation at 72 and 96 h. A modest but significant increase in fungal colony diameter was observed at 48 h in co-culture compared with single culture, whereas subsequent growth reached similar colony sizes in both treatments. By 72 h, the fungal colony had nearly reached the edge of the Petri dish, and complete plate coverage was observed by 96 h regardless of culture condition (Fig. 1B).

**Fig. 1.**
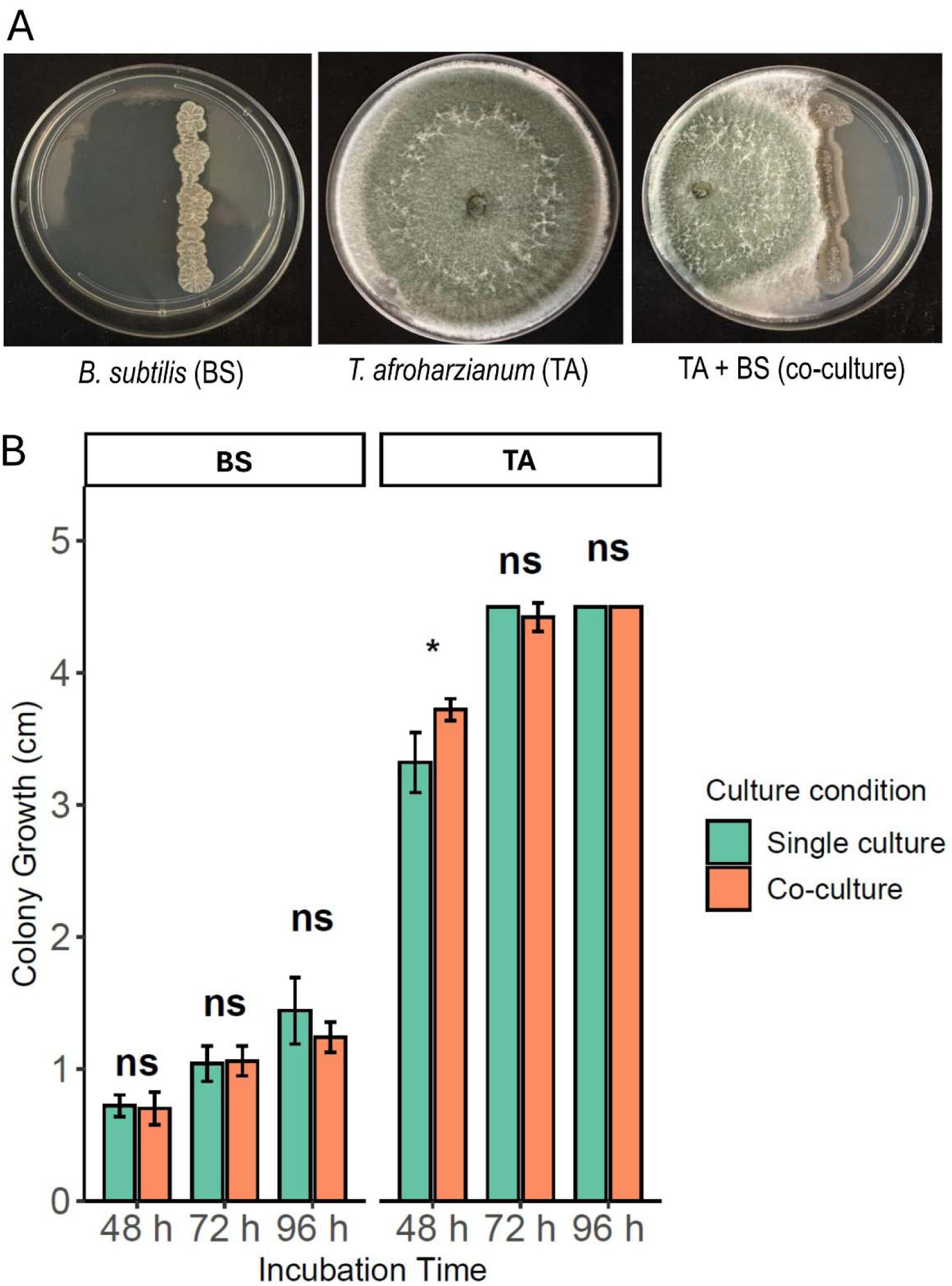
Compatibility of *B. subtilis* (BS) and *T. afroharzianum* (TA) under single and dual-culture conditions. (A) Colony morphology of BS, TA, and their dual culture on half-strength potato dextrose agar (½ PDA). The bacterial inoculum was applied as a single linear streak (approximately 10^8 CFU mL ¹), while a 5-mm mycelial plug of TA was placed on the opposite side of the plate. No inhibition zone or visible changes in colony morphology or sporulation were observed, indicating compatibility between the two microorganisms. (B) Time-course comparison of colony growth in single culture and co-culture at 48, 72, and 96 h. Bars represent the mean ± SD (*n* = 5). Asterisks indicate significant differences between single- and co-culture at each time point based on Student’s t-test (*P* < 0.05), whereas ns denotes no significant difference (*P* ≥ 0.05).

### 3.2. Effects of microbial inoculations on sorghum growth and physiological traits

Microbial inoculation visibly enhanced sorghum growth and root development (Fig. 2A). Sorghum plants inoculated with *B. subtilis*, *T. afroharzianum*, or their co-inoculation (BS+TA) exhibited taller plants and a denser canopy compared to the control plants. Root systems of inoculated plants also appeared larger and more developed than those of the control. Among the inoculated treatments, *B. subtilis* and *T. afroharzianum* co-inoculation displayed the most vigorous overall plant phenotype, with greater shoot and root development (Fig. 2A). PCR analysis confirmed the presence of the respective inoculants in root samples. BS-specific amplification produced the expected 136-bp fragment in the BS and BS+TA treatments, whereas no amplification was detected in the control or TA treatments. Similarly, TA-specific amplification yielded the expected 300-bp fragment in the TA and BS+TA treatments but not in the control or BS treatments (Fig. 2B). The effects of BS, TA, and their combined inoculation (BS+TA) on sorghum growth and physiological performance were evaluated. All microbial treatments significantly increased SPAD chlorophyll content relative to the control, with no significant differences among BS, TA, and BS+TA (Fig. 2C). The maximum quantum efficiency of PSII (Fv/Fm) was significantly higher under TA inoculation, whereas BS and BS+TA showed an intermediate response that was not significantly different from either the control or TA inoculated treatments (Fig. 2D). Similarly, the photosynthetic performance index (Pi_ABS) was significantly greater in all inoculated treatments than in the control, with no significant differences among BS, TA, and BS+TA (Fig. 2E).

**Fig. 2.**
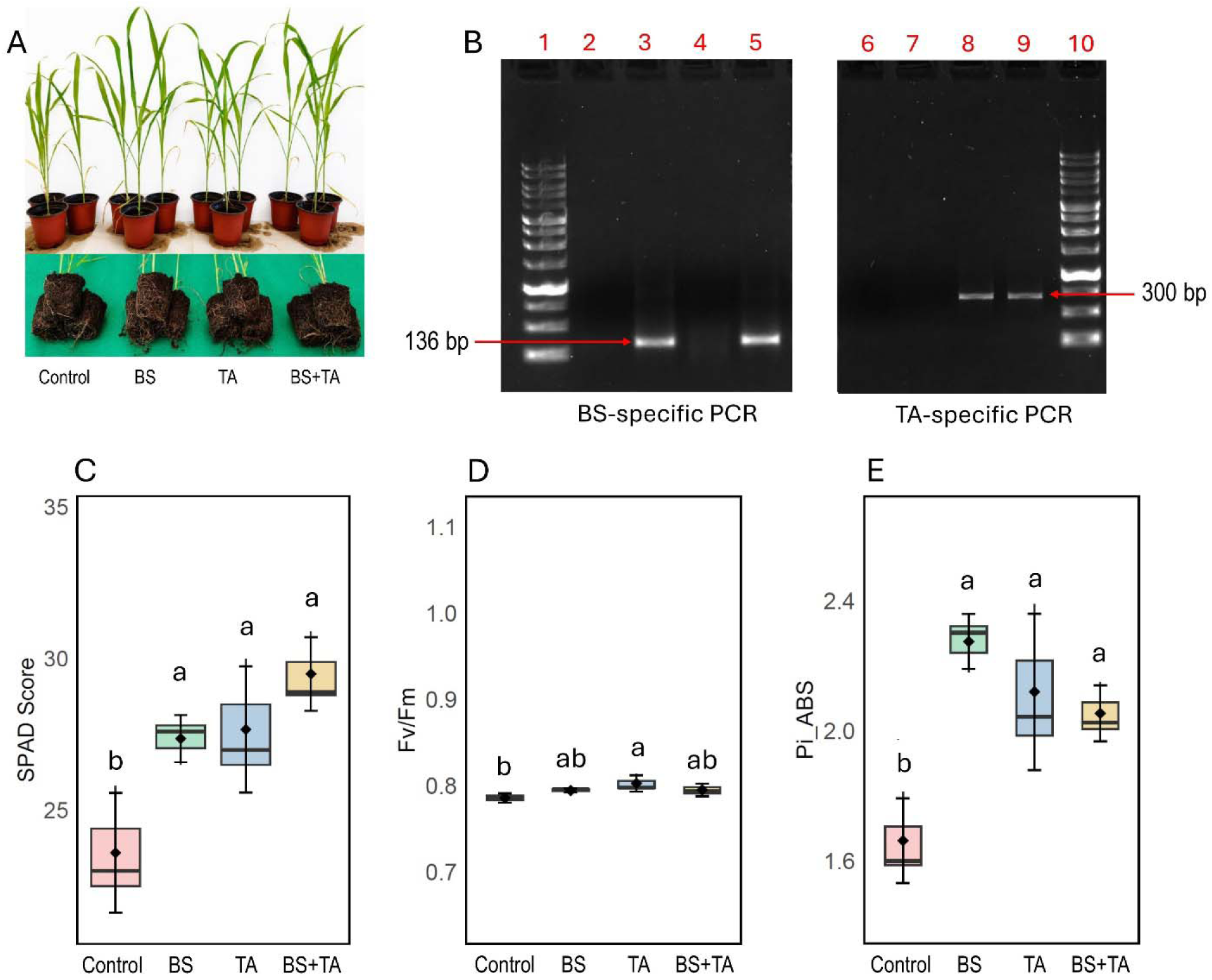
Effects of *B. subtilis* (BS) and *T. afroharzianum* (TA) inoculation on sorghum growth, photosynthetic performance, rhizosphere siderophore production, and PCR-based confirmation of microbial colonization. (A) Phenotype of sorghum plants and root systems under different treatments. (B) PCR confirmation of microbial inoculation using BS-specific (136 bp) and TA-specific (300 bp) primers. Lanes 1 and 10: 1 kb DNA ladder; lanes 2–5 correspond to control, BS, TA, and BS+TA, respectively, for BS-specific PCR; lanes 6–9 correspond to control, BS, TA, and BS+TA, respectively, for TA-specific PCR. (C–E) Box plots showing the effects of treatments on (C) SPAD chlorophyll score, (D) maximum quantum efficiency of PSII (Fv/Fm) and (E) photosynthetic performance index (Pi_ABS). Different letters above the boxes indicate significant differences among treatments according to one-way ANOVA followed by Fisher’s LSD test (*P* < 0.05; *n* = 3 biological replicates).

In this study, shoot height increased significantly in all inoculated treatments relative to the control, while no significant differences were detected among BS, TA, and BS+TA (Fig. 3A). Shoot fresh weight was significantly higher under *B. subtilis–T. afroharzianum* co-inoculation than under the TA- or BS-only treatments, whereas control plants had the lowest shoot fresh weight (Fig. 3B). Root length was significantly greater in BS and BS+TA than in the control, while TA showed an intermediate value that was not significantly different from either group (Fig. 3C). Root fresh weight was significantly increased by BS compared with the control and TA, whereas BS+TA showed an intermediate value that did not differ significantly from BS or TA. In addition, the control exhibited the lowest root fresh weight among all treatments (Fig. 3D). Further, rhizosphere siderophore production also increased significantly following inoculation, with BS, TA, and BS+TA all exhibiting higher values than the control plants (Fig. 3E). In elemental analysis, the inoculation of BS, TA, and BS+TA all exhibited significantly higher Fe concentrations in the root compared with uninoculated controls (Fig. 3F). In leaves, the control plants had the lowest Fe concentration, whereas inoculation with BS, TA, and BS+TA significantly increased Fe concentration. Although the BS+TA consortium exhibited the highest mean leaf Fe concentration, it did not differ significantly from the individual inoculations (Fig. 3F). Both root and leaf N and C concentrations increased following microbial inoculation, with the *B. subtilis* and *T. afroharzianum* consortium showing the highest leaf N and C contents. The control exhibited the lowest root N and C levels, whereas BS, TA, and the BS+TA consortium had significantly higher root N and C (Fig. 3G-H). In contrast, leaf N and C differed among treatments, with the *B. subtilis*-*T. afroharzianum* consortium showing the highest leaf N and C levels relative to the control and other inoculants (Fig. 3G-H).

**Fig. 3.**
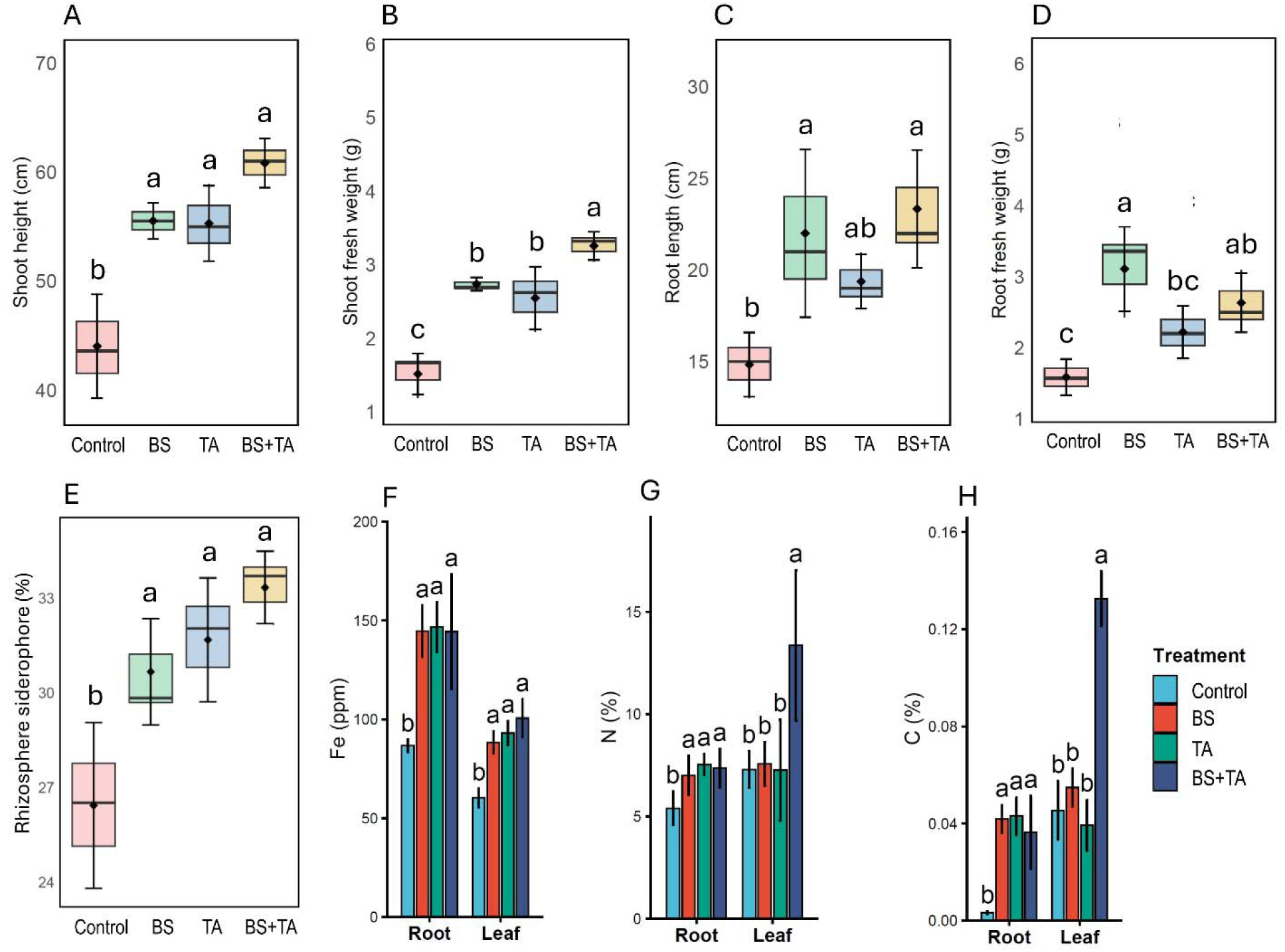
Effects of *B. subtilis* (BS) and *T. afroharzianum* (TA) inoculation on sorghum growth under greenhouse conditions. Box plots show (A) shoot height, (B) shoot fresh weight, (C) root length, (D) root fresh weight, (E) rhizosphere siderophore production and (F, G, H) nutrient status (Fe, N and C) in sorghum inoculated with different inoculants. Different letters above the boxes indicate significant differences among treatments according to one-way ANOVA followed by Fisher’s LSD test (*P* < 0.05; *n* = 3 biological replicates).

### 3.3. Integrated plant performance index

To integrate the effects of microbial inoculation across all measured growth and physiological traits, a Composite Plant Performance Index (CPPI) was calculated based on Z-score standardization (Table 1). The *B. subtilis* and *T. afroharzianum* co-inoculation exhibited the highest CPPI (0.628) and ranked first among all treatments, followed by BS (0.409) and TA (0.270). The control treatment had a negative composite score (−1.308) and ranked fourth. Positive standardized values for most measured traits were observed in the inoculated treatments, whereas the control consistently exhibited negative standardized values across all evaluated parameters (Table 1).

**Table 1.** Composite Plant Performance Index (CPPI) derived from Z-score-standardized growth and physiological traits of sorghum following *B. subtilis* (BS), *T. afroharzianum* (TA), and combined BS+TA inoculation.

| Treatment | SPAD | Fv/Fm | Pi_ABS | Shoot height | Shoot fresh weight | Root length | Root fresh weight | Composite Plant Performance Index (CPPI) | Rank |
| --- | --- | --- | --- | --- | --- | --- | --- | --- | --- |
| Control | -1.176 | -1.032 | -1.364 | -1.418 | -1.425 | -1.383 | -1.355 | -1.308 | 4 |
| BS | -0.154 | -0.010 | 0.921 | 0.231 | 0.318 | 0.684 | 0.874 | 0.409 | 2 |
| TA | 0.326 | 0.973 | 0.345 | 0.197 | 0.047 | -0.060 | 0.060 | 0.270 | 3 |
| BS + TA | 1.004 | 0.069 | 0.098 | 0.990 | 1.059 | 0.759 | 0.421 | 0.628 | 1 |

### 3.4. Response of plants in split-root system

The split-root experiment showed that *B. subtilis* and *T. afroharzianum* (BS+TA) co-inoculation markedly promoted sorghum growth, with the strongest response observed when both root compartments were inoculated (SR3) (Fig. 4B, D–H). Also, PCR analysis confirmed compartment-specific establishment of the microbial inoculants in the split-root system (Fig. 4C). No TA- or BS-specific amplification was detected in either root compartment of SR1 (control/control). In SR2 (control/BS+TA), the expected 300-bp TA-specific and 136-bp BS-specific amplicons were detected only in the inoculated compartment, with no detectable amplification in the uninoculated compartment. In SR3 (BS+TA/BS+TA), both expected amplicons were detected in both root compartments. Further, chlorophyll score and shoot height increased significantly in both SR2 and SR3 compared with SR1, although SR2 and SR3 did not differ significantly from each other (Fig. 4D and 4F). Stem diameter and shoot fresh weight increased progressively across treatments, with SR3 showing the highest value, followed by SR2 and SR1 (Fig. 4E and 4G). Root length increased with BS+TA inoculation compared to controls (Fig. 4H). Within SR1 and SR2, root lengths did not differ significantly between the two root compartments. In contrast, a significant difference (*P* < 0.01) between the two root compartments was observed in SR3 (Fig. 4H).

**Fig. 4.**
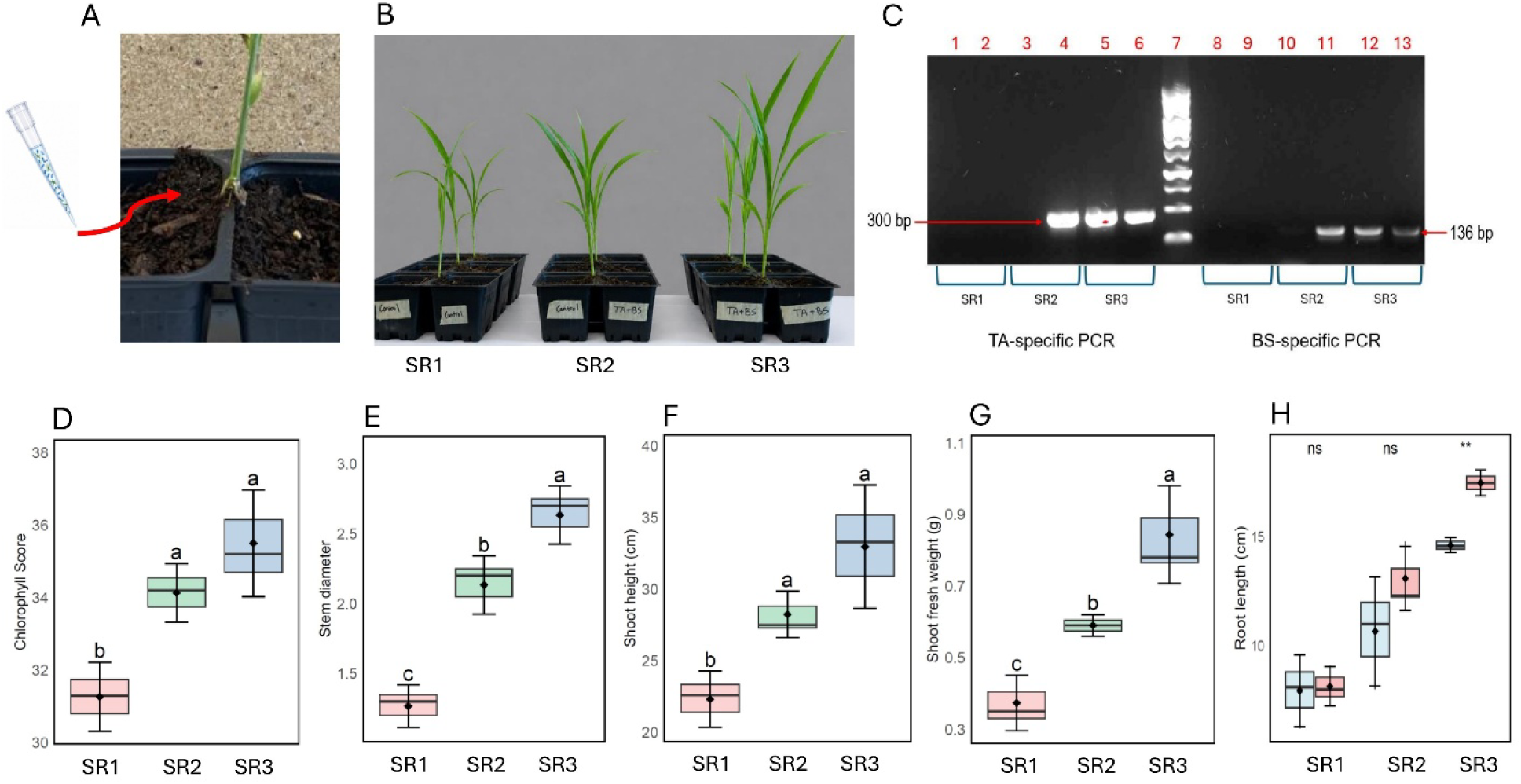
Effects of *B. subtilis* (BS) and *T. afroharzianum* (TA) co-inoculation in a split-root system. Plants were initially grown in sterile soil for 2 weeks and subsequently transferred to a split-root system for an additional 4 weeks. Treatments were SR1 (control/control), SR2 (control/BS+TA), and SR3 (BS+TA/BS+TA). (A) Illustration of microbial inoculation into the root compartment. (B) Phenotype of sorghum plants under the three split-root treatments. (C) Gel electrophoresis image confirming microbial (BS and TA) inoculation in the different root compartments (SR1: control/control, SR2: control/BS+TA, SR3: BS+TA/BS+TA). (D–H) Chlorophyll score, stem diameter, shoot height, shoot fresh weight, and root length, respectively. Different lowercase letters indicate significant differences among treatments, while ns indicates no significant difference; asterisks indicate significant differences between root compartments within each treatment (P < 0.01).

### 3.5. Effects of microbial inoculations on bacterial communities

Non-metric multidimensional scaling (NMDS) based on Bray–Curtis dissimilarity showed the distribution of bacterial community composition among four treatments (Fig. 5A). The NMDS ordination had a stress value of 0.075, indicating a good representation of the community dissimilarities in two dimensions. Samples from the four treatments exhibited modest separation, although some overlap was observed among treatment groups. PERMANOVA detected no significant differences in bacterial community composition among treatments (R² = 0.318, *P* = 0.106). Further, alpha diversity analyses revealed no significant differences among the control, BS, TA, and BS+TA treatments for any of the bacterial diversity indices, including Observed richness and Shannon index (Fig. 5B).

**Fig. 5.**
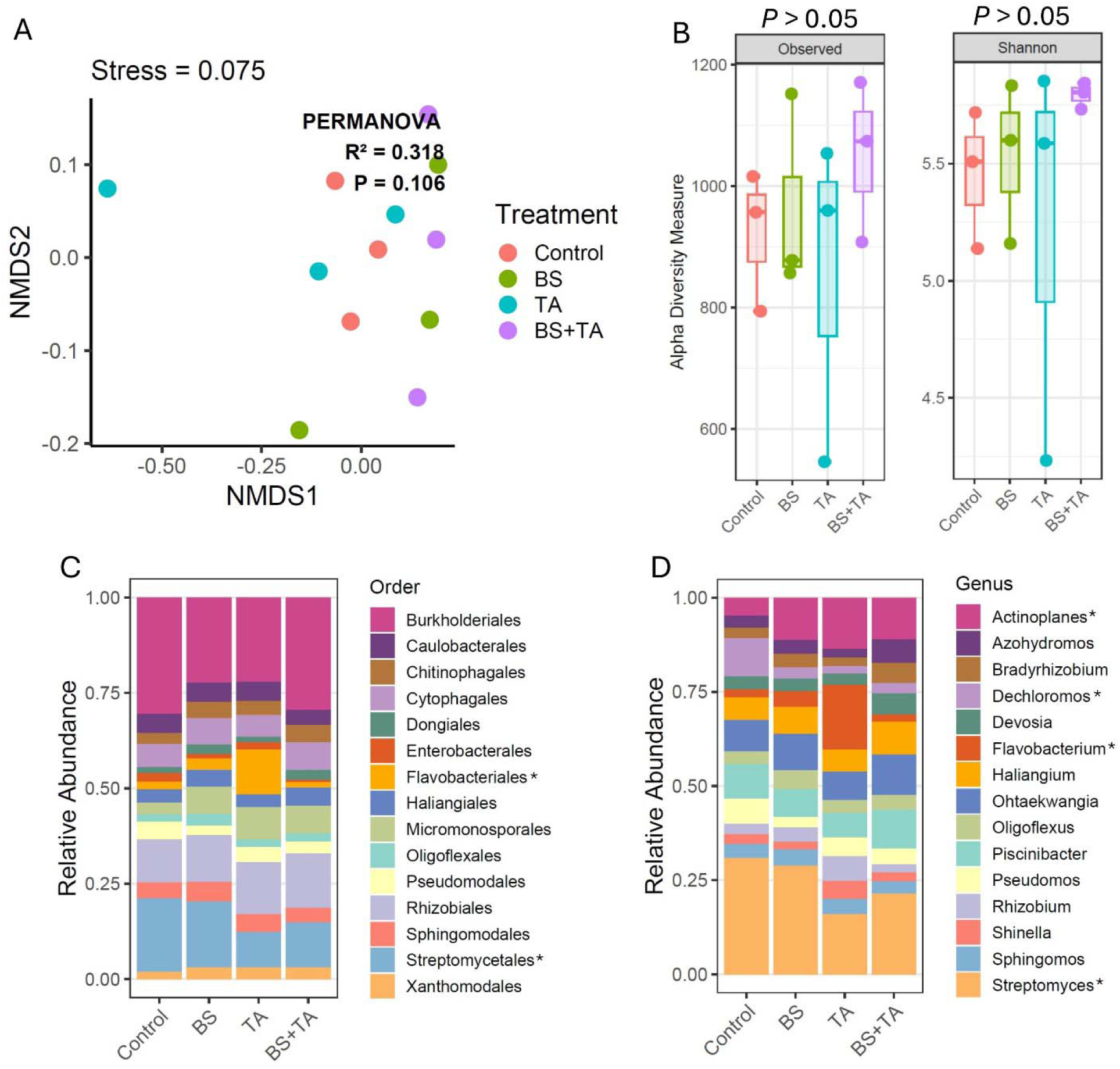
Effects of *B. subtilis* (BS) and *T. afroharzianum* (TA) inoculation on the root bacterial community (16S) of sorghum. (A) Non-metric multidimensional scaling (NMDS) ordination based on Bray–Curtis dissimilarity showing bacterial community composition across different treatments. (B) Alpha diversity of the root bacterial community measured using the Observed richness and Shannon diversity indices. (C) Relative abundance of the dominant bacterial orders across treatments. (D) Relative abundance of the dominant bacterial genera across treatments. Asterisks (*) indicate taxa showing significant differences in relative abundance among treatments (*P* < 0.05). Data represent individual biological replicates (*n* = 3).

At the bacterial order level, the dominant taxa included Xanthomonadales, Streptomycetales, Rhizobiales, Pseudomonadales, Oligoflexales, Micromonosporales, Haliangiales, Flavobacteriales, Enterobacterales, Dongiales, Cytophagales, Chitinophagales, Caulobacterales, and Burkholderiales (Fig. 5C). Among these, Burkholderiales, Flavobacteriales, and Streptomycetales differed significantly among treatments (Fig. 5C). At the bacterial genus level, *Streptomyces* was the most abundant genus across all treatments, followed by *Sphingomonas, Shinella, Rhizobium, Piscinibacter, Oligoflexus, Ohtaekwangia, Haliangium, Flavobacterium, Devosia, Dechloromonas, Bradyrhizobium, Azohydromonas,* and *Actinoplanes* (Fig. 5D). Significant treatment effects were detected for *Actinoplanes, Flavobacterium,* and *Streptomyces* (Fig. 5D).

### 3.6. Effects of microbial inoculation on root-associated fungal communities

In ITS analysis, the NMDS ordination had a stress value of 0.133, indicating an acceptable two-dimensional representation of the community dissimilarities (Fig. 6A). Fungal communities from the four treatments showed substantial overlap with no clear clustering according to treatment. PERMANOVA further indicated that fungal community composition did not differ significantly among treatments (R² = 0.272, *P* = 0.497). Also, no significant differences were detected among treatments for fungal observed richness and Shannon index (Fig. 6B).

**Fig. 6.**
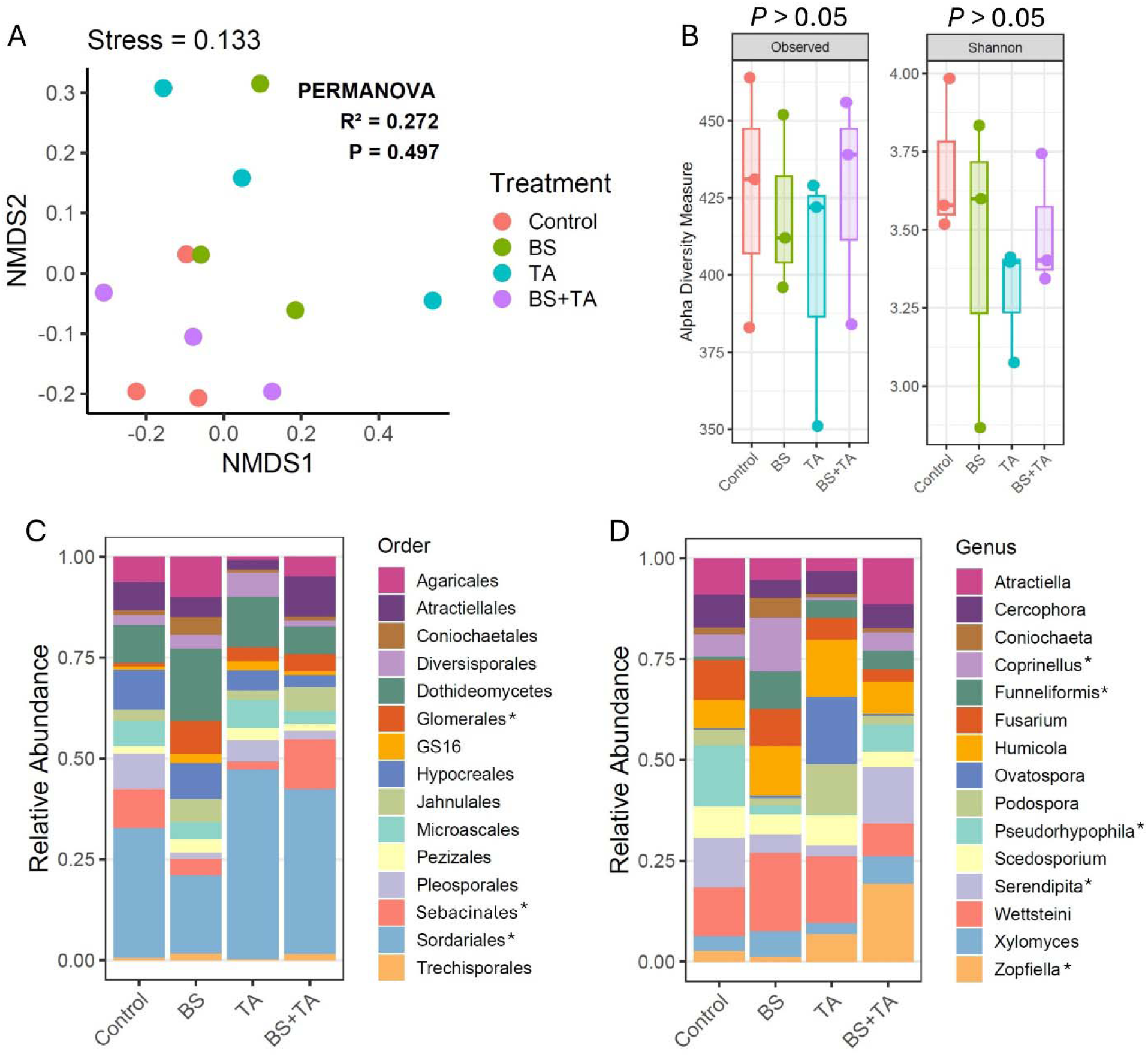
Effects of *B. subtilis* (BS) and *T. afroharzianum* (TA) inoculation on the root fungal community (ITS) of sorghum. (A) Non-metric multidimensional scaling (NMDS) ordination based on Bray–Curtis dissimilarity showing fungal community composition across different treatments. (B) Alpha diversity of the root fungal community measured using the Observed richness and Shannon diversity indices. (C) Relative abundance of the dominant fungal orders across treatments. (D) Relative abundance of the dominant fungal genera across treatments. Asterisks (*) indicate taxa showing significant differences in relative abundance among treatments (*P* < 0.05). Data represent individual biological replicates (*n* = 3).

In fungal analysis, the dominant fungal orders included Trechisporales, Sordariales, Sebacinales, Pleosporales, Pezizales, Microascales, Juhnulales, Hypocreales, GS16, Glomerales, Diversisporales, Coniochaetales, Atractiellales, and Agaricales (Fig. 6C). Among these, only Sordariales, Glomerales and Sebacinales showed a significant difference among treatments (Fig. 6C). At the fungal genus level, the predominant taxa comprised *Zopfiella, Wettsteini, Serendipita, Scedosporium, Pseudohyphopila, Podospora, Ovatospora, Humicola, Fusarium, Funneliformis, Coprinellus, Coniochaeta, Cercophora,* and *Atractiella* (Fig. 6D). Significant differences among treatments were observed for *Coprinellus, Funneliformis, Pseudohyphopila, Serendipita,* and *Zopfiella* (Fig. 6D).

### 3.7. Differential abundance of microbial genera following microbial inoculation

Differential abundance analysis using DESeq2 identified several bacterial amplicon sequence variants (ASVs) that were significantly enriched in response to inoculation relative to the comparison groups (Fig. 7A-C). BS inoculation significantly enriched ASVs assigned to *Actinoplanes* (ASV111, ASV132, ASV176), *Enterobacter* (ASV131 and ASV520) and *Shinella* (ASV473) compared to the controls, whereas *Rhizobium* (ASV237) and *Herbaspirillum* (ASV796) were significantly enriched in the control (Fig. 7A). In contrast, TA inoculation significantly increased the abundance of *Actinoplanes* (ASV132 and ASV176), *Enterobacter* (ASV131), *Shinella* (ASV473), *Pseudomonas* (ASV1641), *Plectobacillus* (ASV1665), and *Flavobacterium* (ASV1666) relative to controls, while *Niastella* (ASV185) was significantly enriched in the control (Fig. 7B). Further, the combined BS+TA treatment significantly enriched *Actinoplanes* (ASV111, ASV132 and ASV176) and *Enterobacter* (ASV131 and ASV520) compared with the control (Fig. 7C). In contrast, *Actinocatus* (ASV179), *Rhizobium* (ASV237), *Ideonella* (ASV277), *Niastella* (ASV185) and *Herbaspirillum* (ASV796) were significantly enriched in control (Fig. 7C).

**Fig. 7.**
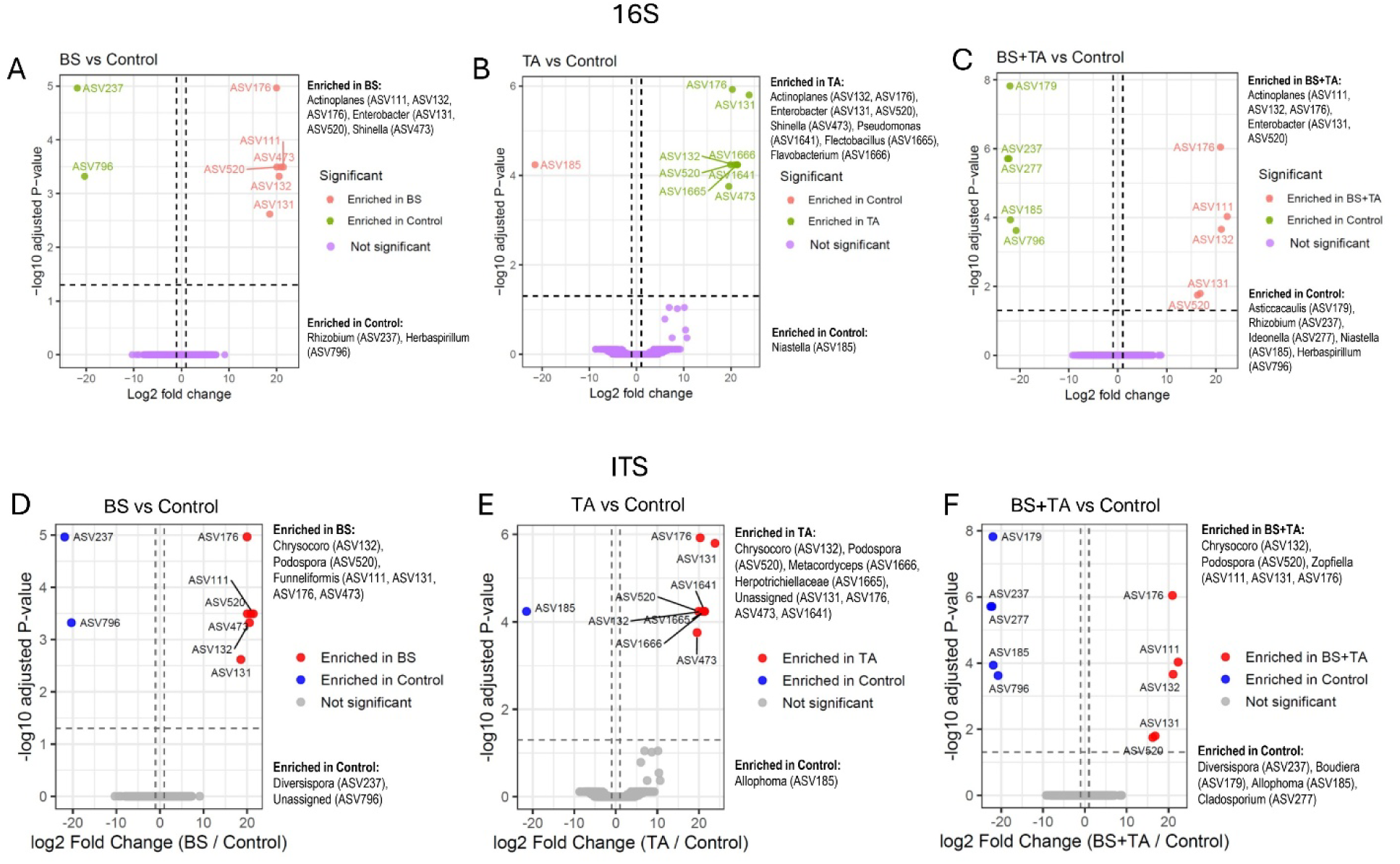
Differentially abundant bacterial and fungal amplicon sequence variants (ASVs) associated with *B. subtilis* (BS), *T. afroharzianum* (TA), and combined BS+TA inoculation in sorghum roots. Volcano plots show differential abundance analyses comparing (A) BS vs Control, (B) TA vs Control, and (C) BS+TA vs Control for the bacterial 16S rRNA dataset, and (D) BS vs Control, (E) TA vs Control, and (F) BS+TA vs Control for the fungal ITS dataset. Each point represents an ASV, with the x-axis indicating log_2_ fold change and the y-axis showing −log_10_ adjusted *P*-value. Red points represent ASVs significantly enriched in the inoculated treatment, blue/green points indicate ASVs significantly enriched in the control, and gray/purple points represent ASVs that were not significantly different. Dashed vertical lines indicate the log_2_ fold-change threshold, and the horizontal dashed line denotes the significance threshold (adjusted *P* < 0.05). Taxonomic identities of significantly enriched ASVs are indicated for each comparison.

DESeq2 analysis identified several fungal amplicon sequence variants (ASVs) that were differentially abundant among the microbial inoculation treatments (Fig. 7D-F). In this study, BS inoculation significantly enriched ASVs assigned to *Chrysocoroo* (ASV132), *Podospora* (ASV520), and unassigned fungal taxa (ASV111, ASV131, ASV176, and ASV473) (Fig. 7D) compared to controls. In contrast, *Diversispora* (ASV237) and an unassigned fungal ASV (ASV796) were significantly enriched in control plants. TA inoculation significantly increased the abundance of *Chrysocoroo* (ASV132), *Podospora* (ASV520), *Metacordyceps* (ASV1666), *Herpotrichiellaceae* (ASV1665), and unassigned fungal ASVs (ASV131, ASV176, ASV473, and ASV1641) relative to controls, whereas *Allophoma* (ASV185) was significantly enriched in the control (Fig. 7E). Furthermore, the combined BS+TA treatment significantly enriched *Chrysocoroo* (ASV132), *Podospora* (ASV520), and unassigned fungal ASVs (ASV111, ASV131, and ASV176) compared with the control (Fig. 7F). Conversely, *Diversispora* (ASV237), *Boudiera* (ASV179), *Allophoma* (ASV185), and *Cladosporium* (ASV277) were significantly enriched in control (Fig. 7F).

### 3.8. Global co-occurrence networks identify treatment-associated bacterial and fungal hub taxa

To identify highly connected microbes associated with each treatment, global bacterial and fungal co-occurrence networks were constructed, and hub taxa were subsequently associated with the treatment in which they exhibited the highest mean relative abundance (Fig. 8A-B). Among bacterial hub taxa associated with control treatment, *Aeromonas, Hydrogenophaga, Sphingobium, Desulfovibrio,* and *Citrifermentans* exhibited the highest degree centrality (Fig. 8A). The BS treatment group was characterized by *Bryobacter*, *Pseudolabrys*, *Salinispira*, *Phenylobacterium*, and Ellin6055, whereas the TA treatment contained *Brevundimonas*, *Shinella*, *Rhizobium*, *Flavobacterium*, and *Delftia* as the principal hub taxa. Among bacterial hub taxa associated with *B. subtilis*-*T. afroharzianum* co-inoculation, *Dongia*, *Lacunisphaera*, *Rhodoplanes*, *Planctomicrobium*, and *Fimbriiglobus* exhibited the highest degree centrality. Distinct fungal hub taxa were also associated with each treatment (Fig. 8B). Among fungal hub taxa, the control treatment was dominated by *Tumularia*, Pseudohyphophila, Lasiosphaeriaceae, *Waitea*, and *Malassezia*. The BS treatment was characterized by *Aspergillus*, Ascobolaceae, *Wettsteinia*, Junewangiaceae, and *Arthrobotrys* as the major hub taxa. Under TA inoculation, *Ectophoma*, *Exserohilum*, *Ovatospora*, *Cephalotrichum*, and *Allophoma* showed the greatest connectivity. In *B. subtilis* and *T. afroharzianum* co-inoculation, Magopthoraceae, *Atractiella*, *Serendipita*, *Zopfiella*, and *Subulicystidium* were identified as the most highly connected fungal hub taxa (Fig. 8B).

**Fig. 8.**
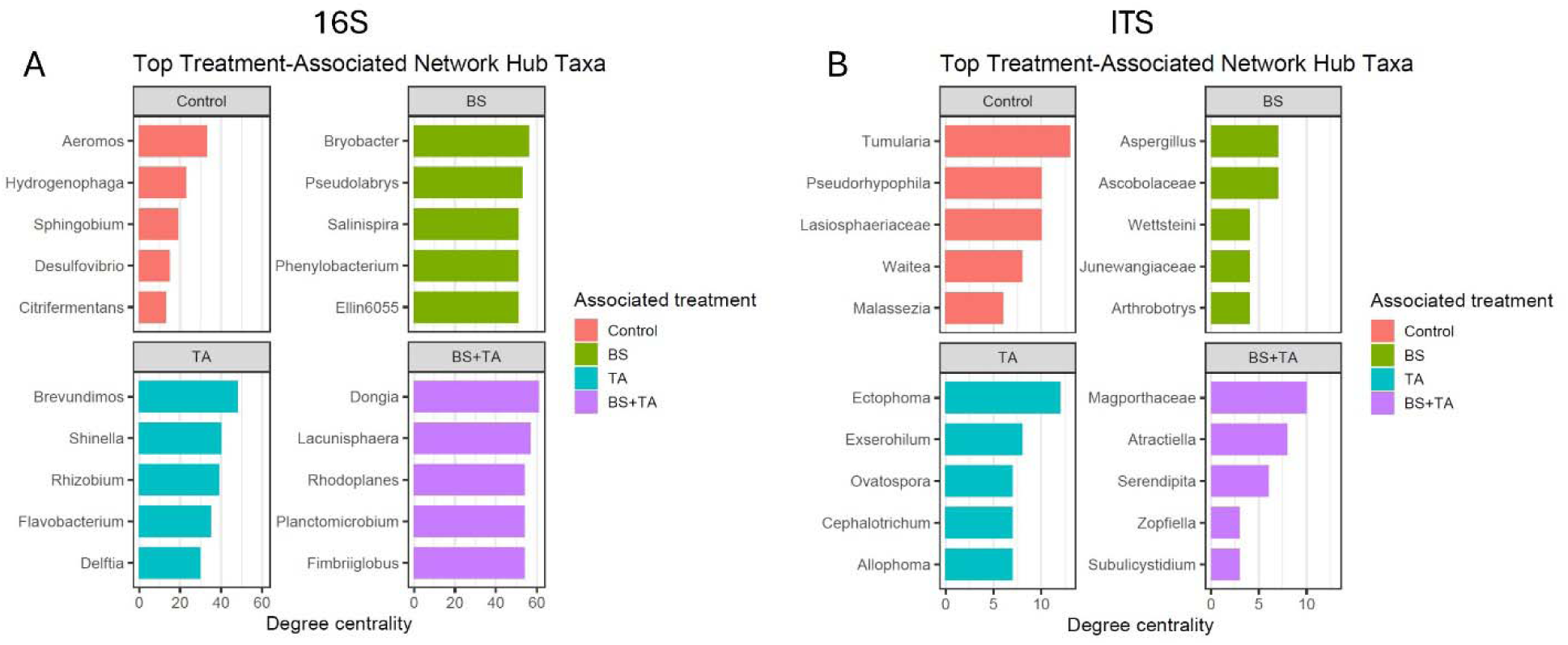
Hub taxa of the sorghum root microbiome identified from co-occurrence network analysis. Top five (A) bacterial (16S rRNA) and (B) fungal hub taxa associated with each treatment (Control, *B. subtilis* [BS], *T. afroharzianum* [TA], and BS+TA), ranked by degree centrality. Hub taxa were identified from global microbial co-occurrence networks and assigned to the treatment in which they exhibited the highest mean relative abundance; higher degree centrality indicates greater connectivity within the microbial community. Bars are colored according to the associated treatment.

### 3.9. Core bacterial and fungal microbiomes of sorghum roots

Core microbiome analysis identified the ten most abundant bacterial and fungal genera meeting the defined prevalence and abundance thresholds. The bacterial core community was dominated by *Streptomyces*, *Rhizobium*, *Pseudomonas*, *Piscinibacter*, *Ohtaekwangia*, *Haliangium*, *Flavobacterium*, *Dechloromonas*, *Azohydromonas*, and *Actinoplanes*, although their relative abundances varied among treatments (Fig. 9A). *Streptomyces* remained one of the most abundant genera across all treatments, while the relative contributions of the other core genera differed between inoculated and non-inoculated plants (Fig. 9A). Similarly, the fungal core microbiome consisted of *Zopfiella*, *Wettsteinia*, *Serendipita*, *Scedosporium*, *Pseudohyphophila*, *Humicola*, *Fusarium*, *Coprinellus*, *Cercophora*, and *Atractiella* (Fig. 9B). These genera were detected in all treatments, with treatment-dependent variation in their relative abundances (Fig. 9B).

**Fig. 9.**
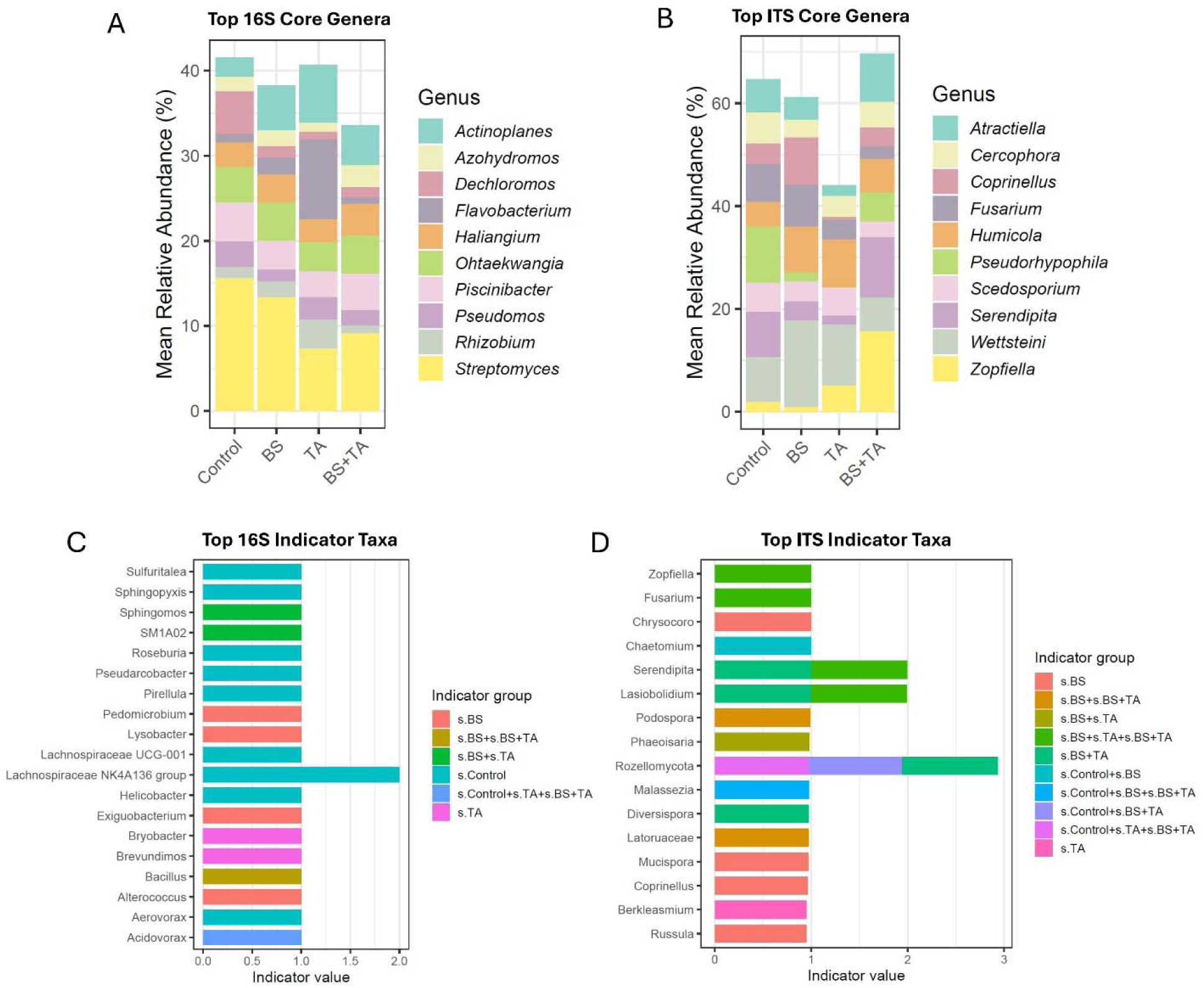
Core and indicator taxa of the sorghum root microbiome associated with *B. subtilis* (BS), *T. afroharzianum* (TA), and combined BS+TA inoculation. Relative abundance of the top 10 core bacterial (A) and fungal (B) genera across the treatments. (C) Top bacterial and (D) Top fungal indicator genera identified by indicator species analysis, with bar lengths representing indicator values and colors denoting the treatment or treatment combination with which each genus was significantly associated. Core taxa represent genera consistently detected across treatments, whereas indicator taxa are significantly associated with specific treatment conditions and may reflect treatment-driven microbial enrichment.

### 3.10. Indicator taxa associated with microbial inoculation

Indicator species analysis identified bacterial and fungal genera that were significantly associated with individual treatments or treatment combinations (Fig. 9C-D). In the bacterial community, indicator genera included *Acidovorax*, *Aerovorax*, *Alterecorax*, *Bacillus*, *Brevundimonas*, *Bryobacter*, *Exiguobacterium*, *Helicobacter*, Lachnospiraceae NK4A136 group, Lachnospiraceae UCG-001, *Lysobacter*, *Pedomicrobium*, *Pirellul*a, *Pseudarcobacter*, *Roseburia*, SM1A02, *Sphingomonas*, *Sphingopyxis*, and *Sulfuritalea* (Fig. 9C). Several taxa were specifically associated with the combined BS+TA treatment, predominantly *Bacillus*, and *Sphingomonas*, whereas Lachnospiraceae NK4A136 group exhibited the highest indicator value and was associated with the control treatment (Fig. 9C). Similarly, fungal indicator analysis identified *Russula*, *Berkleasmium*, *Coprinellus*, *Mucispora*, Laturoaceae, *Diversispora*, *Malassezia*, Rozellomycota, *Phaeoisaria*, *Podospora*, *Lasiobolidium*, *Serendipita*, *Chaetomium*, *Chrysocoroo*, *Fusarium*, and *Zopfiella* as indicator taxa (Fig. 9D). The *B. subtilis* and *T. afroharzianum* consortium was characterized by indicator genera including Rozellomycota, which showed the highest indicator value among all fungal taxa, together with *Serendipita* and *Lasiobolidium*, both of which were associated with inoculated treatment combinations involving *B. subtilis* and *T. afroharzianum* co-inoculation. Additional fungal indicator taxa were associated with the control, BS, TA, or other treatment combinations (Fig. 9D).

### 3.11. Correlation between microbial genera and plant growth traits

Spearman correlation analysis revealed distinct associations between treatment-associated microbial genera and plant growth and physiological traits (Fig. 10A-B). Among the bacterial community, genera associated with the *B. subtilis* and *T. afroharzianum* consortium exhibited the strongest positive correlations with multiple plant performance traits (Fig. 10A). In particular, *Haliangium*, *Azohydromonas*, *Devosia, Dongia, Acidovorax, Pelomonas, Cellulosimicrobium, Quadrisphaera*, *Rhodoplanes*, and *Ramlibacter* showed positive associations with SPAD chlorophyll score, rhizosphere siderophore production, shoot height, shoot fresh weight, and root fresh weight. In contrast, several genera that were most abundant in the control treatment displayed predominantly weak or negative correlations with these traits, while BS- and TA-associated genera exhibited intermediate correlation patterns. A similar trend was observed for the fungal community (Fig. 10B). BS+TA-associated genera, including *Serendipita, Zopfiella, Atractiella, Subulicystidium, Magopthoraceae, Chrysosporium, Oliveonia*, and Rozellomycota, were positively correlated with SPAD chlorophyll score, siderophore production, and growth-related traits. Control-associated fungal genera generally showed weaker or negative correlations, whereas taxa associated with the individual BS or TA treatments displayed mixed positive and negative relationships. Overall, the most consistent positive correlations with plant growth and physiological traits were observed among microbial genera associated with the *B. subtilis* and *T. afroharzianum* consortium (Fig. 10A-B).

**Fig. 10.**
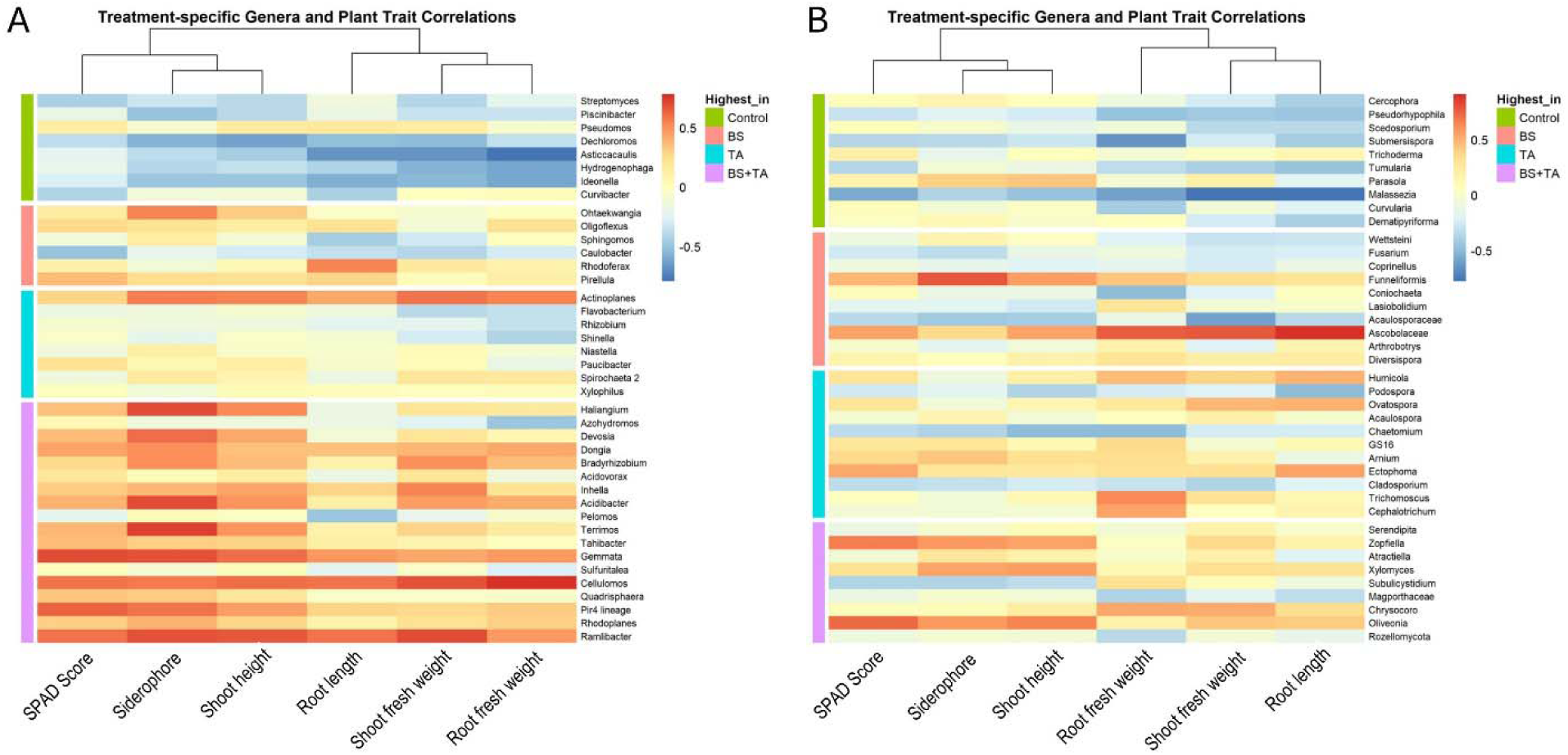
Correlations between dominant microbial genera and plant performance traits under *B. subtilis* (BS), *T. afroharzianum* (TA), and combined BS+TA inoculation. Heatmaps show Spearman correlation coefficients between the relative abundance of treatment-associated microbial genera and plant traits, including SPAD score, rhizosphere siderophore production, shoot height, root length, shoot fresh weight, and root fresh weight. (A) Correlation heatmap for bacterial genera (16S rRNA dataset). (B) Correlation heatmap for fungal genera (ITS dataset). Rows represent microbial genera and columns represent plant traits. Red indicates positive correlations, blue indicates negative correlations, and color intensity reflects the strength of the correlation. Genera are grouped according to the treatment in which they showed the highest relative abundance as indicated by the colored annotation bar. Hierarchical clustering was applied to both genera and plant traits to identify groups with similar correlation patterns.

## 4. Discussion

The present study demonstrates that the *B. subtilis***–***T. afroharzianum* consortium enhances sorghum growth by selectively restructuring the root microbiome rather than altering overall microbial diversity. Integrating plant phenotyping with microbiome profiling, co-occurrence network, core microbiome, indicator species, and plant–microbe correlation analyses revealed how the consortium enriched beneficial microbial taxa, reorganized ecological interactions, and strengthened plant–microbiome associations linked to improved nutrient acquisition and plant performance.

### 4.1. Complementary interactions between *T. afroharzianum* and *B. subtilis* promote sorghum growth

The successful application of microbial consortia requires compatibility among individual microorganisms, allowing them to coexist without antagonistic interactions that could compromise their persistence and beneficial functions (Compant et al., 2019; Santoyo et al., 2021). In this study, the compatibility assay demonstrated that *T. afroharzianum* and *B. subtilis* are compatible under *in vitro* conditions, with no evidence of growth inhibition or morphological alterations during co-cultivation. These results support their simultaneous application as a microbial consortium and establish the experimental basis for subsequent greenhouse experiments investigating their individual and synergistic effects on sorghum performance and rhizosphere microbiome assembly. A recent soybean study showed that co-inoculation with *T. afroharzianum* and *B. subtilis* beneficially altered root and soil microbiomes and concluded that the two inoculants may synergistically promote soybean growth (Rigobelo et al., 2024). The effectiveness of microbial consortia depends not only on the plant growth-promoting capacity of individual strains but also on their ability to coexist and establish stable interactions within the root microbiome (Berg et al., 2020). Many synthetic microbial communities fail because introduced microorganisms compete for similar ecological niches or inhibit one another through antagonistic metabolites (Toju et al., 2018). Therefore, the successful establishment of both *T. afroharzianum* and *B. subtilis* in sorghum roots indicates that both microorganisms simultaneously colonized the root system, suggesting compatibility between the bacterial and fungal partners. Such compatibility is increasingly recognized as a key factor governing the persistence and effectiveness of microbial inoculants under greenhouse and field conditions (Panek et al., 2026).

The superior performance of sorghum under the *B. subtilis*–*T. afroharzianum* co-inoculation is therefore likely attributable to the complementary functional roles of the two microorganisms rather than redundancy in their modes of action. Increasing evidence indicates that bacterial–fungal consortia often outperform individual inoculants because microorganisms occupying different ecological niches collectively provide a broader range of ecosystem services than either partner alone (Berg et al., 2020; Woo et al., 2023). *Bacillus* rapidly colonizes the rhizosphere through biofilm formation and produces numerous metabolites involved in plant growth promotion, including indole-3-acetic acid (IAA), volatile organic compounds, lipopeptides, siderophores, and phosphate-solubilizing enzymes (Hashem et al., 2019; Blake et al., 2021; Mahapatra et al., 2022). In contrast, *Trichoderma* spp., including *T. afroharzianum,* establish intimate associations with root tissues, where they secrete auxin-like compounds, cellulases, and siderophores that stimulate root branching and Fe uptake in plants (Kabir et al., 2024; Kabir and Bennetzen, 2024). Photosynthesis is closely linked to plant mineral nutrition, particularly Fe, which serves as an essential cofactor for chlorophyll biosynthesis and photosynthetic electron transport (Marschner, 2012). The coordinated increase in Fe concentrations in root and leaf, rhizosphere siderophore production, and photosynthetic parameters following microbial inoculation suggests that enhanced microbial Fe mobilization improved plant performance. Besides facilitating Fe acquisition, siderophore-producing microorganisms can promote the establishment of other beneficial microorganisms possessing compatible Fe acquisition systems (Ahmed & Holmström, 2014). Furthermore, the greater leaf N and C levels observed with the combined BS+TA treatment is consistent with previous reports demonstrating complementary or synergistic interactions between bacterial and fungal inoculants that enhance nutrient translocation and photosynthetic carbon assimilation more effectively than either microorganism alone (Colla et al., 2015; Rouphael & Colla, 2020). Further, the greater N and C in plant tissues, particularly in the leaves of sorghum inoculated with the BS–TA consortium, indicates that co-inoculation enhanced nutrient acquisition and assimilation by the host plant. Increased C levels in leaves may further reflect enhanced photosynthetic performance and carbohydrate production (Thompson et al., 2017). This is commonly associated with improved mineral nutrition and microbial-mediated stimulation of plant metabolism (Su et al., 2024). Carbon and nitrogen metabolism are tightly interconnected, whereas increased C-fixation supplies the carbon skeletons required for amino acid biosynthesis (Zheng, 2009). Beneficial microorganisms may therefore improve overall metabolic efficiency by simultaneously promoting carbon assimilation and nitrogen acquisition (Singh et al., 2022). Overall, the coordinated enhancement of Fe, N, and C contents indicates that the BS–TA consortium improves nutrient uptake and metabolic efficiency more effectively than either microorganism alone.

To further provide an integrated assessment, our CPPI analysis ranked the *B. subtilis*–*T. afroharzianum* consortium as the highest-performing treatment, followed by *B. subtilis* and *T. afroharzianum*. Unlike individual physiological measurements, which evaluate specific aspects of plant function, composite indices integrate multiple traits into a single metric and therefore provide a more holistic assessment of overall plant performance (Poorter et al., 2012). The superior CPPI of the *T. afroharzianum* and *B. subtilis* treatment supports the concept that combining bacterial and fungal inoculants can generate complementary effects across multiple plant functions rather than improving only a single trait. Also, microbial consortia with complementary ecological strategies may provide a more effective approach for developing next-generation microbial biofertilizers for sustainable sorghum production.

### 4.2. Localized root inoculation drives BS+TA-mediated growth promotion

Beneficial microbes are known to initiate such local-to-systemic communication. For example, Trichoderma-derived signals are perceived locally by roots, where they alter physiological processes such as Fe acquisition, while subsequently generated signals can travel to distal tissues and modify shoot responses (Martínez-Medina et al., 2017). The split-root experiment provides evidence that the growth-promoting effect of *B. subtilis*–*T. afroharzianum* consortium is primarily initiated through localized interactions at the inoculated roots. Our PCR analysis confirms that *B. subtilis* and *T. afroharzianum* remained localized to the inoculated root compartments, with no detectable microbial transfer to the uninoculated side. Inoculation of only one root compartment (SR2) improved some whole-plant traits, including chlorophyll status and shoot height, but bilateral inoculation (SR3) generally produced the strongest growth response. These findings suggest that local recognition and colonization of the consortium can initiate root-derived signals that subsequently influence shoot physiology and growth. However, inoculation of both root compartments (SR3) generally resulted in the greatest growth promotion, indicating that the magnitude of the response depends on the extent of direct root–microbe interaction.

This pattern is consistent with our previous study in sorghum showing that *T. afroharzianum* T22 acts predominantly through local root signaling, with direct fungal colonization inducing transcriptional and endophytic microbiome changes associated with growth promotion (Kabir et al., 2024). In contrast, our recent split-root study with *B. subtilis* in garden pea demonstrated that locally applied bacteria can also generate responses beyond the inoculated root region, supporting a systemic component of Bacillus-mediated signaling (Kabir et al., 2026). Thus, in the present consortium, local perception of *B. subtilis* and *T. afroharzianum* may represent the primary trigger, while downstream root-to-shoot signaling allows these localized interactions to translate into whole-plant growth responses. The stronger phenotype observed when both root compartments received *B. subtilis*–*T. afroharzianum* consortium further suggests that systemic signaling alone does not fully substitute for direct microbial contact. Instead, greater root coverage likely expands the sites of microbial perception, nutrient mobilization, and microbiome modulation, thereby amplifying the resulting whole-plant response. Collectively, the split-root results support a model in which localized *B. subtilis*–*T. afroharzianum* co-inoculation is required to initiate growth-promoting signals, with increasing spatial colonization of the root system strengthening these signals and maximizing whole-plant growth promotion.

### 4.3. Microbial inoculation selectively restructures the sorghum root microbiome without altering overall diversity

Although neither alpha diversity nor beta diversity differed significantly among treatments, several dominant bacterial and fungal taxa exhibited clear shifts in relative abundance following inoculation. This pattern suggests that inoculated *B. subtilis* and *T. afroharzianum* primarily influenced community composition through selective recruitment or suppression of particular taxa rather than causing wholesale changes in microbial diversity. Similar observations have been reported, where beneficial inoculants modify the abundance of specific microbial groups while preserving overall community structure (Compant et al., 2019; Toju et al., 2018). Further, root-associated microbial communities are generally assembled through strong host selection, in which plants recruit microorganisms capable of utilizing root exudates while excluding poorly adapted taxa (Bulgarelli et al., 2013; Fitzpatrick et al., 2020). Consequently, introducing beneficial microorganisms does not necessarily increase species’ richness but instead alters the relative abundance of microorganisms already adapted to the root environment (de Vries et al., 2020).

The differential enrichment of *Actinoplanes* and *Enterobacter* indicates that *B. subtilis*-*T. afroharzianum* consortium recruited functionally distinct microbial groups that may contribute to nutrient cycling and plant growth promotion. It suggests that co-inoculation reshaped the existing community through ecological filtering rather than wholesale replacement of resident microorganisms. *Actinoplanes*, which was significantly enriched across all inoculation treatments, is known to produce extracellular enzymes and bioactive metabolites that contribute to organic matter decomposition (Barka et al., 2016). Similarly, *Enterobacter*, *Pseudomonas*, and *Flavobacterium*, which were enriched primarily by *T. afroharzianum*, include strains capable of biological nitrogen metabolism, siderophore production, and improved nutrient acquisition (Compant et al., 2019; Mahapatra et al., 2022). On the other hand, the *B. subtilis***–***T. afroharzianum* co-inoculation primarily enriched *Podospora*, a genus containing plant growth-promoting fungi. *P. bulbillosa* has been shown to enhance drought and salinity tolerance in tomato by strengthening antioxidant defenses, promoting osmolyte accumulation, and regulating stress-responsive genes (Kazerooni et al., 2022). These observations indicate that *B. subtilis* and *T. afroharzianum*, particularly when applied as a consortium, selectively favored microbial taxa possessing complementary functional roles to sorghum plants. Therefore, selective microbiome modulation appears to be a key feature of effective microbial inoculants for sorghum growth.

### 4.4. Ecological reorganization of the root microbiome following combined inoculation

Microbial communities are ecological networks in which community stability depends on both species composition and interactions among microorganisms rather than on individual taxa alone (Kodera et al., 2022; Philippot et al., 2021). In the present study, the persistence of dominant bacterial and fungal genera, especially *Streptomyces* and *Serendipita*, indicates that the *B. subtilis***–***T. afroharzianum* consortium did not replace the resident microbiome but instead modified the relative abundance and ecological roles of indigenous microorganisms. *Streptomyces* is involved in the decomposition of complex organic substrates, nutrient mineralization, and suppression of soilborne pathogens (Viaene et al., 2016). Further, *Serendipita* is a beneficial root endophytic fungus that promotes nutrient acquisition, modulates phytohormone signaling, and induces antioxidant defenses, thereby increasing tolerance to both biotic and abiotic stresses (Rong et al., 2023; Saleem et al., 2022). Unlike dominant taxa, hub microorganisms frequently regulate microbial interactions by connecting otherwise weakly linked components of ecological networks (Agler et al., 2016). The *B. subtilis*–*T. afroharzianum* consortium formed a distinct co-occurrence network with *Rhodoplanes*, *Serendipita*, and *Zopfiella* as hub taxa. This indicates that microbial inoculation reshaped ecological interactions beyond changes in community composition. Although the ecological role of *Rhodoplanes* remains poorly understood, *Serendipita* is well known to help plants cope with abiotic stresses through mutualistic interactions with host plants (Saleem et al., 2022; Shekhawat et al., 2021). In addition, *Zopfiella* species are saprotrophic fungi involved in the decomposition of organic matter and nutrient turnover (Mahongnao et al., 2024; Floudas et al., 2020). Thus, the emergence of distinct hub taxa following *B. subtilis*–*T. afroharzianum* co-inoculation suggests a reorganization of microbial community connectivity and potential ecological associations, although the functional roles of these hubs require experimental validation.

Indicator species analysis further identified distinct microbial groups associated with inoculation, with *Bacillus* and *Serendipita* among the indicator taxa associated with treatment combinations involving BS+TA co-inoculation. Indicator taxa differ from core microorganisms because they exhibit strong specificity and fidelity to particular environmental conditions, making them useful biomarkers of ecological change (De Cáceres & Legendre, 2009). The enrichment of *Bacillus* as an indicator taxon suggests a beneficial rhizosphere microbiome, as *Bacillus* spp. promote nutrient acquisition and stress tolerance (Tsotetsi et al., 2022). Similar plant-beneficial functions have also been reported for Serendipita (Saleem et al., 2022). Thus, the simultaneous occurrence of distinct hub taxa and indicator species suggests that co-inoculation altered both community composition and ecological interactions within the sorghum microbiome. These findings support the possibility that successful microbial inoculation may be accompanied by changes in microbial community organization rather than changes in overall diversity alone.

### 4.5. Plant–microbiome associations identify candidate microbial taxa linked to improved sorghum performance

To further link microbiome assembly with plant performance, we examined associations between microbial taxa and plant growth and physiological traits. Although correlation does not establish causality, it helps identify microorganisms potentially associated with improved plant performance (Agler et al., 2016). The predominance of positive correlations between inoculation-associated microorganisms and sorghum performance suggests that the *B. subtilis–T. afroharzianum* consortium reshaped the microbiome while strengthening beneficial associations with the host. For example, *Devosia* is recognized for its ability to colonize roots, fix atmospheric nitrogen in association with certain hosts, and improve nutrient availability under nutrient-limited conditions (Rivas et al., 2003). The fungal community exhibited a similar trend, with Serendipita positively associated with plant traits consistent with its established plant-beneficial functions.

Interestingly, several genera that exhibited positive correlations with plant traits were not necessarily the most abundant within the root microbiome. This suggests that microorganisms need not be highly abundant to be associated with plant performance, particularly when they occupy highly connected positions within microbial communities. These coordinated microbial responses are consistent with the observed improvements in plant traits and microbiome restructuring, suggesting that the consortium promoted coordinated enhancement of sorghum performance. Overall, the enrichment and positive associations of key bacterial and fungal genera indicate that the superior performance of the *B. subtilis*–*T. afroharzianum* co-inoculation was accompanied by coordinated changes in root microbial composition and plant–microbiome associations. These beneficial taxa represent promising targets for next-generation bioinoculants and provide a foundation for microbiome-assisted improvement of sorghum. However, the verification of the ecological roles of the identified taxa will require targeted approaches such as synthetic community (SynCom) experiments, microbial isolation, and genome-resolved metagenomics to determine how these microorganisms contribute to nutrient cycling and plant growth.

## 5. Conclusion

The *B. subtilis–T. afroharzianum* consortium enhanced sorghum growth and physiological performance, increased tissue carbon levels, and promoted rhizosphere siderophore production, consistent with improved plant nutrient status. Split-root analysis demonstrated that bilateral root co-inoculation was required to maximize whole-plant growth benefits. Rather than altering overall microbial diversity, the consortium selectively restructured the root microbiome by enriching beneficial taxa (*Actinoplanes*, *Enterobacter*, and *Podospora*). Also, *Streptomyces* and *Serendipita* persisted as prominent members of the core microbiome, and BS+TA-associated hub and indicator taxa revealed selective changes in microbial community organization (Fig. 11). These findings provide a microbiome-informed basis for developing bacterial–fungal consortia as next-generation bioinoculants for sustainable sorghum production.

**Fig. 11.**
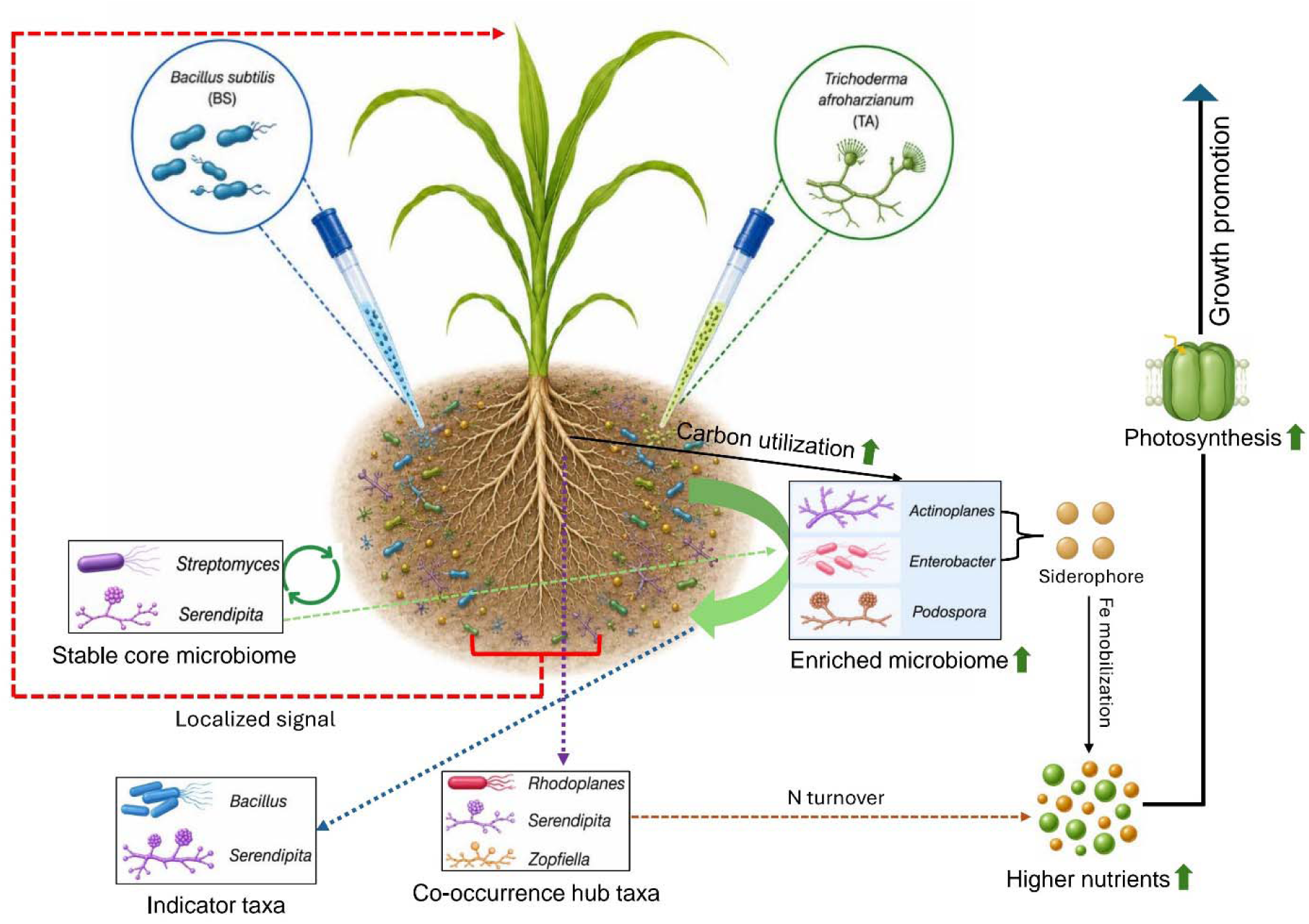
Proposed conceptual model illustrating how the *B. subtilis* (BS)**–***T. afroharzianum* (TA) consortium improves growth in sorghum. Localized root signaling initiated by *B. subtilis***-***T. afroharzianum* co-inoculation may coordinate host physiological responses and microbiome restructuring, ultimately enhancing nutrient acquisition, photosynthesis, and plant growth. Co-inoculation was associated with increased plant carbon status and selective enrichment of beneficial microorganisms, including *Actinoplanes*, the siderophore-producing *Enterobacter*, and *Podospora*. The persistence of potentially beneficial core taxa, including *Streptomyces* and *Serendipita*, may contribute to sustained rhizosphere functioning. Indicator species and co-occurrence network analyses further identified inoculation-associated taxa, with *Rhodoplanes, Serendipita,* and *Zopfiella* among prominent hub taxa potentially contributing to microbial community connectivity and organization

## Acknowledgements

The authors thank LC Sciences (Houston, TX, USA) for performing amplicon sequencing and for providing high-quality sequencing data. We also acknowledge the technical support provided throughout the sequencing process.

## Data availability statement

The raw amplicon sequencing data generated in this study are publicly available in the NCBI BioProject database under accession numbers: PRJNA1495209 (16S) and PRJNA1495211 (ITS) (https://www.ncbi.nlm.nih.gov/bioproject/).

## References

1. Agler, M. T., Ruhe, J., Kroll, S., Morhenn, C., Kim, S. T., Weigel, D., & Kemen, E. M. (2016). Microbial hub taxa link host and abiotic factors to plant microbiome variation. PLoS Biology, 14(1), e1002352.

2. Ahmed, E., & Holmström, S. J. M. (2014). Siderophores in environmental research: Roles and applications. Microbial Biotechnology, 7(3), 196–208.

3. Alexander, D. B., & Zuberer, D. A. (1991). Use of chrome azurol S reagents to evaluate siderophore production by rhizosphere bacteria. Biology and Fertility of Soils, 12, 39–45.

4. Berendsen, R. L., Pieterse, C. M. J., & Bakker, P. A. H. M. (2012). The rhizosphere microbiome and plant health. Trends in Plant Science, 17(8), 478–486.

5. Berg, G., Rybakova, D., Fischer, D., Cernava, T., Vergès, M. C. C., Charles, T., Chen, X., Cocolin, L., Eversole, K., Corral, G. H., Kazou, M., Kinkel, L., Lange, L., Lima, N., Loy, A., Macklin, J. A., Maguin, E., Mauchline, T., McClure, R., … Schloter, M. (2020). Microbiome definition re-visited: Old concepts and new challenges. Microbiome, 8, 103.

6. Blake, C., Christensen, M. N., & Kovács, Á. T. (2021). Molecular aspects of plant growth promotion and protection by *Bacillus subtilis*. Molecular Plant-Microbe Interactions, 34, 15–25.

7. Bulgarelli, D., Schlaeppi, K., Spaepen, S., van Themaat, E. V. L., & Schulze-Lefert, P. (2013). Structure and functions of the bacterial microbiota of plants. Annual Review of Plant Biology, 64, 807–838.

8. Callahan, B. J., McMurdie, P. J., Rosen, M. J., Han, A. W., Johnson, A. J. & Holmes, S. P. (2016) DADA2: High-resolution sample inference from Illumina amplicon data. Nature Methods, 13(7), 581–583.

9. Clarke J. D. (2009). Cetyltrimethyl ammonium bromide (CTAB) DNA miniprep for plant DNA isolation. Cold Spring Harbor protocols, 2009(3), pdb.prot5177.

10. Compant, S., Samad, A., Faist, H., & Sessitsch, A. (2019). A review on the plant microbiome: Ecology, functions, and emerging trends in microbial application. Journal of Advanced Research, 19, 29–37.

11. Contreras-Cornejo, H. A., et al. (2024). Mechanisms for plant growth promotion activated by *Trichoderma* spp. Applied Soil Ecology, 196, 105309.

12. De Cáceres, M., & Legendre, P. (2009). Associations between species and groups of sites: Indices and statistical inference. Ecology, 90(12), 3566–3574.

13. de Vries, F. T., Griffiths, R. I., Knight, C. G., Nicolitch, O., & Williams, A. (2020). Harnessing rhizosphere microbiomes for drought-resilient crop production. Science, 368(6488), 270–274. 10.1126/science.aaz5192

14. Fitzpatrick, C. R., Salas-González, I., Conway, J. M., et al. (2020). The plant microbiome: From ecology to reductionism and beyond. Annual Review of Microbiology, 74, 81–100.

15. Floudas, D., Bentzer, J., Ahrén, D., Johansson, T., Persson, P., & Tunlid, A. (2020). Uncovering the hidden diversity of litter-decomposition mechanisms in mushroom-forming fungi. The ISME journal, 14(8), 2046–2059.

16. Gastélum, G., Gómez-Gil, B., Olmedo-Álvarez, G., & Rocha, J. (2025). Harnessing emergent properties of microbial consortia for Agriculture: Assembly of the Xilonen SynCom. Biofilm, 9, 100284.

17. Hartman, K., van der Heijden, M. G. A., Roussely-Provent, V., Walser, J.-C., & Schlaeppi, K. (2018). Deciphering composition and function of the root microbiome of a legume plant. Microbiome, 6, 2. 10.1186/s40168-017-0381-2

18. Hashem, A., Tabassum, B., & Abd_Allah, E. F. (2019). *Bacillus subtilis*: A plant-growth-promoting rhizobacterium that also impacts biotic stress. Saudi Journal of Biological Sciences, 26, 1291–1297.

19. Jacoby, R., Peukert, M., Succurro, A., Koprivova, A., & Kopriva, S. (2017). The role of soil microorganisms in plant mineral nutrition—Current knowledge and future directions. Frontiers in Plant Science, 8, 1617.

20. Kabir, A. H., Thapa, A., Hasan, M. R., & Mostofa, M. (2026). *Bacillus subtilis* reprograms host transcriptome and rhizosphere microbiome via systemic signaling to confer alkaline stress tolerance in garden pea. bioRxiv. 10.1101/2025.09.13.676035

21. Kabir, A. H., Thapa, A., Hasan, M. R., & Parvej, M. R. (2024). Local signal from *Trichoderma afroharzianum* T22 induces host transcriptome and endophytic microbiome leading to growth promotion in sorghum. Journal of experimental botany, 75(22), 7107–7126.

22. Kabir, A. H., & Bennetzen, J. L. (2024). Molecular insights into the mutualism that induces iron deficiency tolerance in sorghum inoculated with *Trichoderma harzianum*. Microbiological research, 281, 127630.

23. Kazerooni, E. A., Maharachchikumbura, S. S. N., Al-Sadi, A. M., Rashid, U., Kang, S. M., & Lee, I. J. (2022). *Actinomucor elegans* and *Podospora bulbillosa* Positively Improves Endurance to Water Deficit and Salinity Stresses in Tomato Plants. Journal of fungi (Basel, Switzerland), 8(8), 785.

24. Kodera, S. M., Das, P., Gilbert, J. A., & Lutz, H. L. (2022). Conceptual strategies for characterizing interactions in microbial communities. iScience, 25(2), 103775.

25. Mahapatra, S., Yadav, R., & Ramakrishna, W. (2022). *Bacillus subtilis* impact on plant growth, soil health and environment. Journal of Applied Microbiology, 132, 3543–3562.

26. Mahongnao, S., Sharma, P., & Nanda, S. (2024). Characterization of fungal microbiome structure in leaf litter compost through metagenomic profiling for harnessing the bio-organic fertilizer potential. 3 Biotech, 14(9), 191.

27. Marschner, P. (2012). Marschner’s mineral nutrition of higher plants (3rd ed.). Academic Press.

28. Martínez-Medina, A., Van Wees, S. C. M., & Pieterse, C. M. J. (2017). Airborne signals from Trichoderma fungi stimulate iron uptake responses in roots resulting in priming of jasmonic acid-dependent defences in shoots of *Arabidopsis thaliana* and *Solanum lycopersicum*. Plant, cell & environment, 40(11), 2691–2705.

29. McMurdie, P. J., & Holmes, S. (2013). phyloseq: an R package for reproducible interactive analysis and graphics of microbiome census data. PloS one, 8(4), e61217.

30. Neilands, J. B. (1995). Siderophores: Structure and function of microbial iron transport compounds. Journal of Biological Chemistry, 270(45), 26723–26726.

31. Panek, J., Gryta, A., Maj, W., Mącik, M., Oszust, K., Pertile, G., Pylak, M., Siegieda, D., Hallama, M., Hatano, R., Kandeler, E., Pathan, S. I., Pietramellara, G., Malusa, E., Weber, J., Turnau, K., Różalska, S., & Frąc, M. (2026). Plant-soil-microbiome interactions: mechanisms, advances, and challenges in sustainable agriculture and healthy agroecosystems. Frontiers in microbiology, 17, 1762743.

32. Philippot, L., Griffiths, B. S., & Langenheder, S. (2021). Microbial Community Resilience across Ecosystems and Multiple Disturbances. Microbiology and molecular biology reviews : MMBR, 85(2), e00026–20.

33. Poorter, H., Niklas, K. J., Reich, P. B., Oleksyn, J., Poot, P., & Mommer, L. (2012). Biomass allocation to leaves, stems and roots: Meta-analyses of interspecific variation and environmental control. New Phytologist, 193(1), 30–50.

34. Quast, C., Pruesse, E., Yilmaz, P., Gerken, J., Schweer, T., Yarza, P., Peplies, J., & Glöckner, F. O. (2013). The SILVA ribosomal RNA gene database project: improved data processing and web-based tools. Nucleic acids research, 41(Database issue), D590–D596.

35. Reid, T. E., et al. (2024). Local signal from *Trichoderma afroharzianum* T22 induces host transcriptome and endophytic microbiome leading to growth promotion in sorghum. Journal of Experimental Botany, 75, 7107–7126.

36. Rigobelo, E. C., de Andrade, L. A., Santos, C. H. B., Frezarin, E. T., Sales, L. R., de Carvalho, L. A. L., Guariz Pinheiro, D., Nicodemo, D., Babalola, O. O., Verdi, M. C. Q., Mondin, M., & Desoignies, N. (2024). Effects of Trichoderma harzianum and Bacillus subtilis on the root and soil microbiomes of the soybean plant INTACTA RR2 PRO™. Frontiers in plant science, 15, 1403160.

37. Rivas, R., Willems, A., Subba-Rao, N. S., Mateos, P. F., Dazzo, F. B., Kroppenstedt, R. M., Martínez-Molina, E., Gillis, M., & Velázquez, E. (2003). *Devosia* species associated with plants: Taxonomy and ecological significance. International Journal of Systematic and Evolutionary Microbiology, 53, 1893–1900.

38. Rong, Z. Y., Lei, A. Q., Wu, Q. S., Srivastava, A. K., Hashem, A., Abd Allah, E. F., Kuča, K., & Yang, T. (2023). Serendipita indica promotes P acquisition and growth in tea seedlings under P deficit conditions by increasing cytokinins and indoleacetic acid and phosphate transporter gene expression. Frontiers in plant science, 14, 1146182.

39. Rooney, W. L., Blumenthal, J., Bean, B., & Mullet, J. E. (2007). Designing sorghum as a dedicated bioenergy feedstock. Biofuels, Bioproducts and Biorefining, 1(2), 147–157.

40. Sahib, M. R., et al. (2020). Rhizobacterial species richness improves sorghum growth and soil nutrient availability. Scientific Reports, 10, 14856.

41. Saiz-Fernández, I., Černý, M., Skalák, J., & Brzobohatý, B. (2021). Split-root systems: detailed methodology, alternative applications, and implications at leaf proteome level. Plant methods, 17(1), 7.

42. Saleem, S., Sekara, A., & Pokluda, R. (2022). Serendipita indica-A Review from Agricultural Point of View. Plants (Basel, Switzerland), 11(24), 3417.

43. Santoyo, G., Guzmán-Guzmán, P., Parra-Cota, F. I., Santos-Villalobos, S. D. L., Orozco-Mosqueda, M. D. C., & Glick, B. R. (2021). Plant growth stimulation by microbial consortia. Agronomy, 11(2), 219.

44. Shade, A., & Handelsman, J. (2012). Beyond the Venn diagram: The hunt for a core microbiome. Environmental Microbiology, 14(1), 4–12.

45. Shekhawat, P. K., Jangir, P., Bishnoi, A., Roy, S., Ram, H., & Soni, P. (2021). Serendipita indica: Harnessing its versatile potential for food and nutritional security. Physiological and Molecular Plant Pathology, 116, 101708. 10.1016/j.pmpp.2021.101708

46. Singh, S. K., Wu, X., Shao, C., & Zhang, H. (2022). Microbial enhancement of plant nutrient acquisition. Stress biology, 2(1), 3.

47. Su, F., Zhao, B., Dhondt-Cordelier, S., & Vaillant-Gaveau, N. (2024). Plant-Growth-Promoting Rhizobacteria Modulate Carbohydrate Metabolism in Connection with Host Plant Defense Mechanism. International journal of molecular sciences, 25(3), 1465.

48. Thompson, M., Gamage, D., Hirotsu, N., Martin, A., & Seneweera, S. (2017). Effects of Elevated Carbon Dioxide on Photosynthesis and Carbon Partitioning: A Perspective on Root Sugar Sensing and Hormonal Crosstalk. Frontiers in physiology, 8, 578.

49. Timmusk, S., Behers, L., Muthoni, J., Muraya, A., & Aronsson, A.-C. (2017). Perspectives and challenges of microbial application for crop improvement. Frontiers in Plant Science, 8, 49. 10.3389/fpls.2017.00049

50. Toju, H., Peay, K. G., Yamamichi, M., Narisawa, K., Hiruma, K., Naito, K., Fukuda, S., Ushio, M., Nakaoka, S., & Onoda, Y. (2018). Core microbiomes for sustainable agroecosystems. Nature Plants, 4(5), 247–257. 10.1038/s41477-018-0139-4

51. Tsotetsi, T., Nephali, L., Malebe, M., & Tugizimana, F. (2022). Bacillus for Plant Growth Promotion and Stress Resilience: What Have We Learned?. Plants (Basel, Switzerland), 11(19), 2482.

52. Vejan, P., Abdullah, R., Khadiran, T., Ismail, S., & Nasrulhaq Boyce, A. (2016). Role of plant growth promoting rhizobacteria in agricultural sustainability—A review. Molecules, 21(5), 573.

53. Viaene, T., Langendries, S., Beirinckx, S., Maes, M., & Goormachtig, S. (2016). *Streptomyces* as a plant’s best friend? FEMS Microbiology Ecology, 92(8), fiw119. 10.1093/femsec/fiw119

54. Zheng Z. L. (2009). Carbon and nitrogen nutrient balance signaling in plants. Plant signaling & behavior, 4(7), 584–591.

